# The *Trichoplax* microbiome: the simplest animal lives in an intimate symbiosis with two intracellular bacteria

**DOI:** 10.1101/568287

**Authors:** Harald R. Gruber-Vodicka, Nikolaus Leisch, Manuel Kleiner, Tjorven Hinzke, Manuel Liebeke, Margaret McFall-Ngai, Michael G. Hadfield, Nicole Dubilier

## Abstract

Placozoa is an enigmatic phylum of simple, microscopic, marine metazoans. Although intracellular bacteria have been found in all members of this phylum, almost nothing is known about their identity, location and interactions with their host. We used metagenomic and metatranscriptomic sequencing of single host individuals, plus metaproteomic and imaging analyses, to show that the placozoan *Trichoplax* H2 lives in symbiosis with two intracellular bacteria. One symbiont forms a new genus in the Midichloriaceae (Rickettsiales) and has a genomic repertoire similar to that of rickettsial parasites, but does not appear to express key genes for energy parasitism. Correlative microscopy and 3-D electron tomography revealed that this symbiont resides in an unusual location, the rough endoplasmic reticulum of its host’s internal fiber cells. The second symbiont belongs to the Margulisbacteria, a phylum without cultured representatives and not known to form intracellular associations. This symbiont lives in the ventral epithelial cells of *Trichoplax*, likely metabolizes algal lipids digested by its host, and has the capacity to supplement the placozoan’s nutrition. Our study shows that even the simplest animals known have evolved highly specific and intimate associations with symbiotic, intracellular bacteria, and highlights that symbioses with microorganisms are a basal trait of animal life.

## Main

Placozoa is a phylum of marine invertebrates at the base of the animal tree, whose members are considered the simplest animals known. These minute, flat and amoeba-like animals of only 0.2 – 2 mm diameter have no mouth or gut, no organs, no muscle or nerve cells, and arose relatively soon after the transition from unicellular to multicellular organisms. Placozoans can be easily cultured, and are considered key models for understanding metazoan evolution, developmental biology and tissue formation^1-4^. Electron microscopy studies as early as the 1970s revealed the presence of intracellular bacteria in these animals^5-8^. Remarkably, nearly five decades later, still only very little is known about the biology of these symbionts and their interactions with their hosts.

The phylum Placozoa occurs in temperate to tropical oceans, and consists of two genera, *Trichoplax* and *Hoilungia*, and at least 19 cryptic species, called haplotypes^8-10^. These benthic animals feed on algae and bacterial biofilms by external digestion and subsequent uptake via the ventral epithelium^11,12^. All known placozoans consist of six morphologically differentiated cell types that are organized in three layers^5,8,13,14^. The thick ventral epidermis consists of ciliated epithelial cells, in which glandular and lipophilic cells are irregularly interspersed. The thin dorsal epidermis consists of ciliated epithelial cells in which crystal cells occasionally occur. An internal meshwork of fiber cells, sandwiched between the two epidermal layers, connects the ventral and dorsal body wall^14^. Intracellular symbionts were first described from these fiber cells^5,7,14^. The bacteria were present in all seven haplotypes examined, independent of sampling site or time, and were hypothesized to reside in the lumen of the rough endoplasmic reticulum (rER)^5,7,8,14^. Persistent and stable residence of a bacterium in the rER of a host would be remarkable, as the vast majority of intracellular symbionts live in the cytoplasm or vacuoles, and the few known exceptions inhabit the nucleus or mitochondria^15-17^.

Sequencing projects of placozoan genomes consistently yielded rickettsial and other bacterial sequences ^8,18,19^. However, as thousands of host individuals were pooled for these analyses, it was neither clear if these bacterial sequences originated from contaminants or symbionts nor if they were consistently present in all host individuals. Our recent advances in sequencing both the metagenome and metatranscriptome of single host individuals with DNA and RNA yields as low as 0.5 ng, together with correlative imaging analyses, allowed us to explore the patterns, structure, and function of the placozoan symbiosis at the individual and cellular level. We focused on the *Trichoplax* haplotype H2, previously reported to host two bacterial morphotypes^7^. To characterize the microbiome of this placozoan, we employed a combination of metagenomic, metatranscriptomic, and metaproteomic analyses together with fluorescence in situ hybridization, and 3-D reconstruction based on serial electron microscopy tomography.

## Results and Discussion

### The *Trichoplax* H2 microbiome is dominated by two bacterial symbionts

We isolated a placozoan H2 haplotype lineage from a seawater tank at the Kewalo Marine Laboratory, University of Hawai‘i at Mānoa, Honolulu, Hawai‘i (Supplementary Fig. 1). To characterize the microbiome of this *Trichoplax* H2, we combined highly sensitive DNA and RNA extraction and library preparation protocols, to sequence the metagenomes and metatranscriptomes of microscopic single individuals that have an estimated biovolume of 0.02 µl and from which we could isolate 0.5 to 4 ng of nucleic acids (n=5). Based on 16S ribosomal RNA (rRNA) gene reads, all five individuals had similar microbial communities, consisting only of Alphaproteobacteria, Gammaproteobacteria and Flavobacteria as well as a member of the Margulisbacteria, a recently characterized phylum with no cultured representatives^20,21^ (Supplementary Fig. 2). Only two taxa from these bacterial phyla were highly abundant in all five host individuals (Supplementary Table 1).

The first, and most abundant 16S rRNA phylotype was an Alphaproteobacterium from the family Midichloriaceae (Rickettsiales)^22^ (Fig. 1a). Midichloriaceae are obligate intracellular, often pathogenic, bacteria found in protists and animals, including humans^23^. In 16S rRNA analyses, the *Trichoplax* H2 midichloriacean phylotype formed a new lineage that clustered with sequences recovered from diverse invertebrate hosts, including the cnidarian *Hydra*, the Pacific oyster (*Crassostrea gigas*) and the Japanese spiky sea cucumber (*Apostichopus japonicus*), as well as sequences from subsurface sediment samples (98.4% – 99.4% identity).

**Figure 1.**
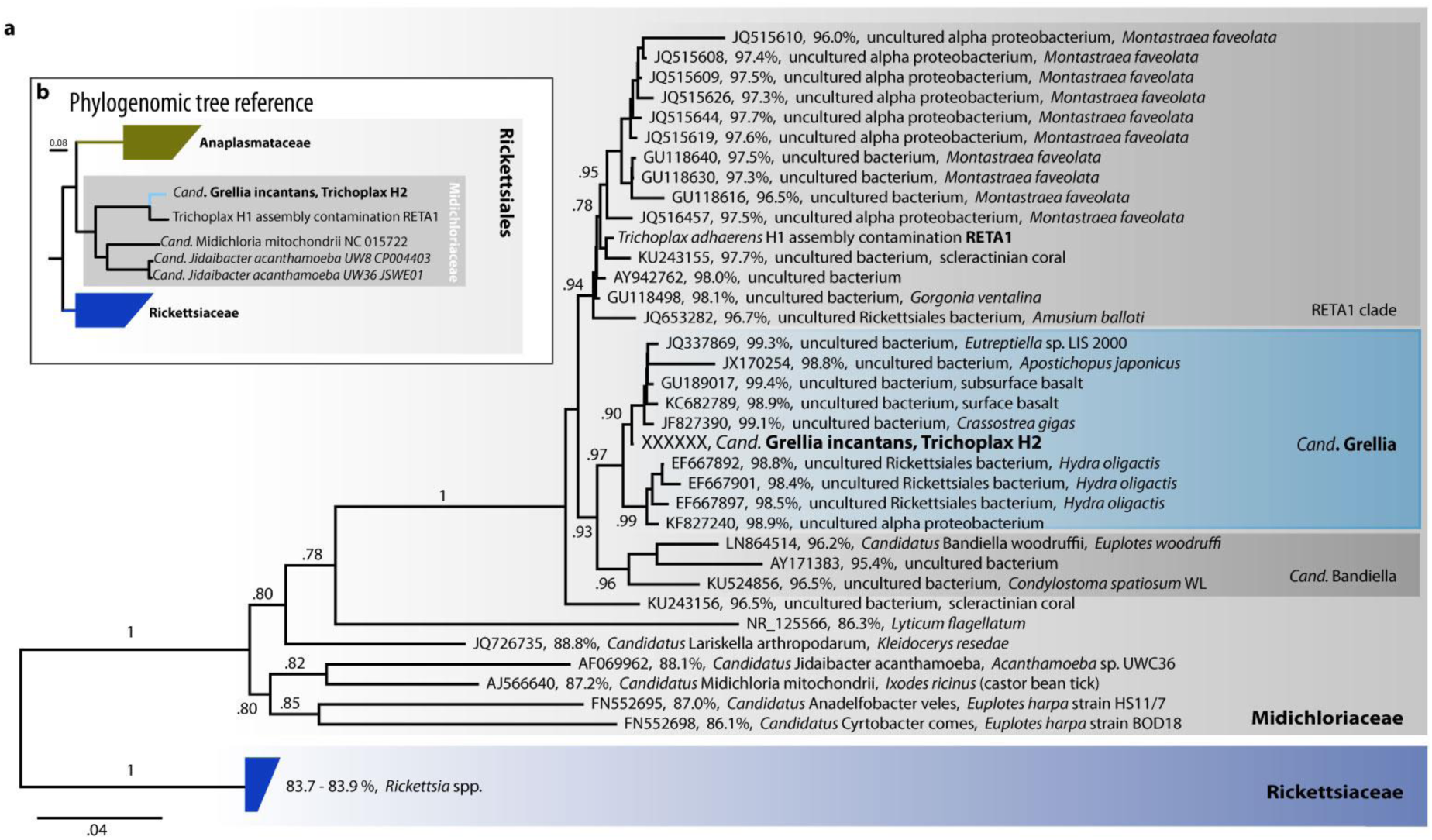
*Cand*. Grellia incantans represents a novel genus in Midichloriaceae (Rickettsiales) Bootstrap support values below 0.5 are not shown. Scale bars indicate substitutions per site. **a**, 16S rRNA tree of G. incantans and related Midichloriaceae; for each sequence, the accession number, the % identity to G. incantans, and the published taxonomic names and hosts, where available, are indicated. **b**, Phylogenomic reconstruction using 43 conserved marker genes based on metagenome assembled genomes and reference genomes.

We propose the *Candidatus* taxon Grellia incantans for this midichloriacean phylotype, based on tree topology and 16S rRNA gene identities of 95.4 – 96.5 % to the closest characterized genus *Cand*. Bandiella^24,25^ (G. incantans from here on; see Supplementary Note 1 for detailed description and etymology). In the sequence data from the *Trichoplax adhaerens* haplotype H1 genome project^2^, a 600-bp fragment of the 16S rRNA gene of a rickettsial phylotype was detected^19^. The 16S rRNA sequence of this phylotype was 98.3% identical to that of G. incantans. Based on tree topology, the *Trichoplax* H1 phylotype belongs neither to the genus Grellia nor to *Cand*. Bandiella, but to a separate, yet undescribed genus we gave the working name RETA1 here (Fig. 1a).

We employed a metagenomic binning strategy based on assembly graph analysis^26^ that enabled us to recover the complete 1.26 Mb bacterial chromosome of G. incantans (Supplementary Note 1). We used the genome of G. incantans to BLAST-search the *Trichoplax* H1 genome for related sequences, and assembled a partial genome of a rickettsial bacterium (RETA1) that was highly similar to the set of rickettsial contigs found in the *Trichoplax* H1 genome by Driscoll *et al*.^19^. Phylogenomic analyses of the G. incantans genome, the RETA1 draft genome and selected Rickettsiales corroborated our 16S rRNA gene analyses and placed the *Trichoplax* H1 and H2 symbionts in the Midichloriaceae. G. incantans was phylogenetically distinct from the *Trichoplax* H1 RETA1 and, based on amino acid sequence identity, these two symbionts belong to two separate genera (Fig. 1a and 1b, Supplementary Note 1)^27,28^.

The second most abundant and consistently present bacterial taxon in the *Trichoplax* H2 metagenomes belonged to the Margulisbacteria, a phylum without isolated representatives that forms the sister clade to Cyanobacteriota^21,29-31^. No 16S rRNA sequences with > 90% identity to this bacterial taxon were found in public sequence databases, indicating a novel group at the genus or even family level. We propose a new *Candidatus* taxon Ruthmannia eludens for this bacterium (R. eludens from here on; see Supplementary Note 2 for detailed description and etymology).

Using metagenomics binning, we recovered a 1.51 Mb metagenome assembled genome for R. eludens with an average GC content of 37%. Our phylogenomic analyses confirmed our 16S rRNA gene results and placed R. eludens in the Margulisbacteria^21,29^ (Fig. 2). Three classes of Margulisbacteria are currently characterized in the Genome Taxonomy Database (GTDB: http://gtdb.ecogenomic.org), WOR-1, GWF2-35-9, and ZB3 (Marinamargulisbacteria)^21,32^, while a fourth class is known from termites (Termititenax)^33^.

**Figure 2.**
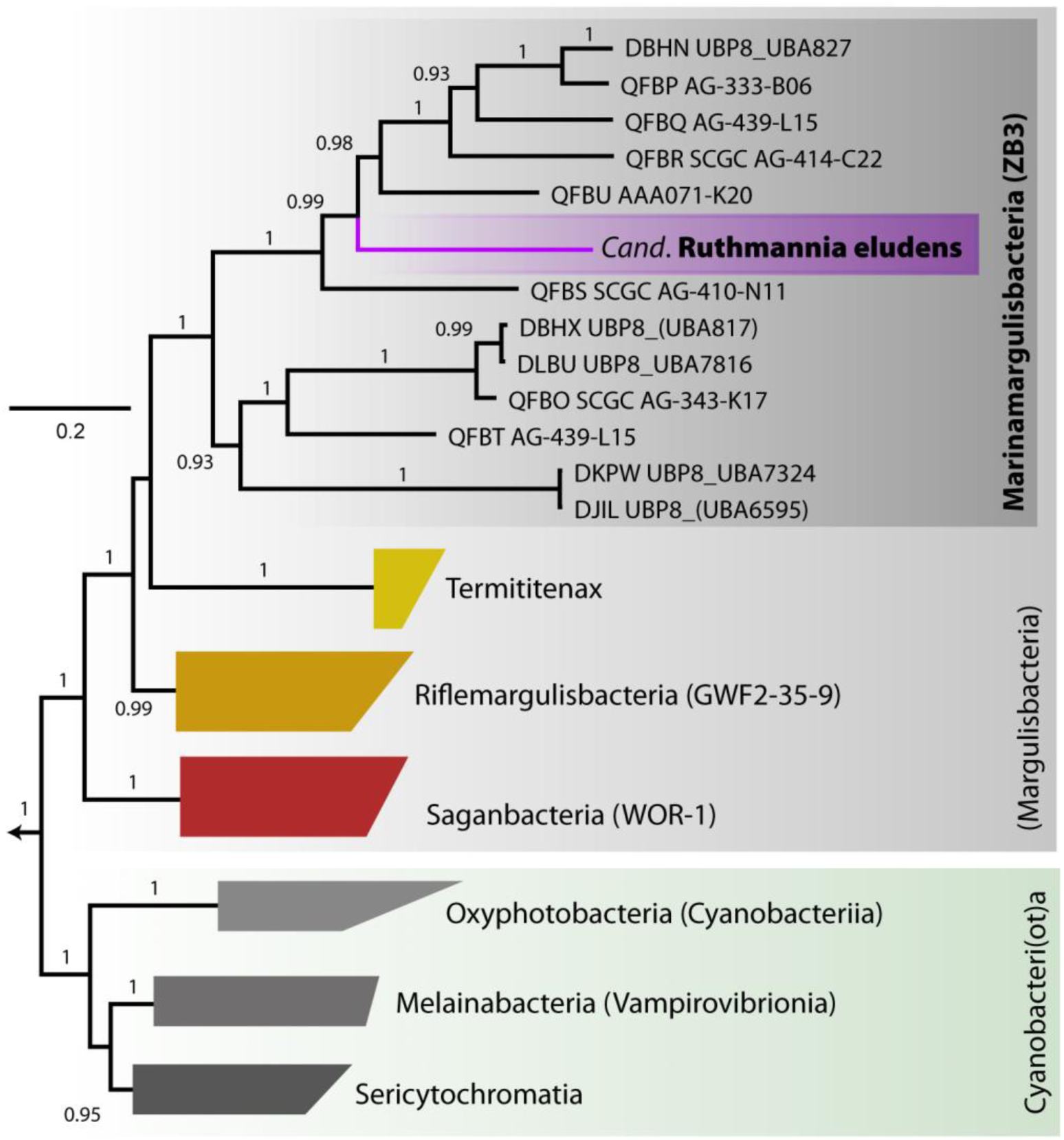
*Cand*. Ruthmannia eludens is a Marinamargulisbacterium (Margulisbacteria). Phylogenomic reconstruction using 43 conserved marker genes based on metagenome assembled genomes, single cell amplified genomes and reference genomes. Phylum-level classification follows GTDB. Taxon names in GTDB are indicated in parenthesis where available. Scale bar indicates substitutions per site. Boot strap support values below 0.5 are not shown.

R. eludens belongs to the Marinamargulisbacteria and was distantly related to several single-cell amplified genomes and metagenome-assembled genomes from oceanic samples^21^ (Fig. 2). Marinamargulisbacteria are aquatic bacteria that occur worldwide in a wide range of water and sediment samples, and are only known from sequence-based studies; draft genomes have been recovered only from marine pelagic samples (Fig. 2). Only seven single-cell amplified genomes and five metagenome-assembled genomes are available for Marinamargulisbacteria, with recovered drafts of 0.5 – 2.0 Mb, and all genomes are classified as medium to low quality ^34^. Our binning strategy led to the recovery of a complete bacterial chromosome from the Margulisbacteria with a genome size of 1.5 Mb (Supplementary Note 2).

### Both symbionts are intracellular, spatially segregated and specific to host cell type

To link the bacterial sequences to their morphotypes and visualize the distribution of the two symbionts in *Trichoplax*, we used fluorescence *in situ* hybridization (FISH) with probes specific to the two symbionts as well as a general probe for Bacteria (Supplementary Table 2). To overcome the high autofluorescence of the host and improve the signal to noise ratio, we modified the standard FISH protocol and used double and quadruple labeled probes, combined with highly sensitive microscopy (Supplementary Fig. 3). No bacteria except the two symbionts G. incantans and R. eludens were detected in all placozoan individuals examined (Fig. 3a; Supplementary Fig. 4). G. incantans was thin and rod-shaped, with a maximum length of 1.2 μm and width of 0.20 to 0.30 μm (Fig. 3a and Supplementary Figs. 4). In contrast, R. eludens had a wider and stouter rod-shaped morphology with a similar maximum length but a width of 0.33 to 0.47 μm (Fig. 3a and Supplementary Figs. 4) (for details see Supplementary Notes 1 and 2).

**Figure 3.**
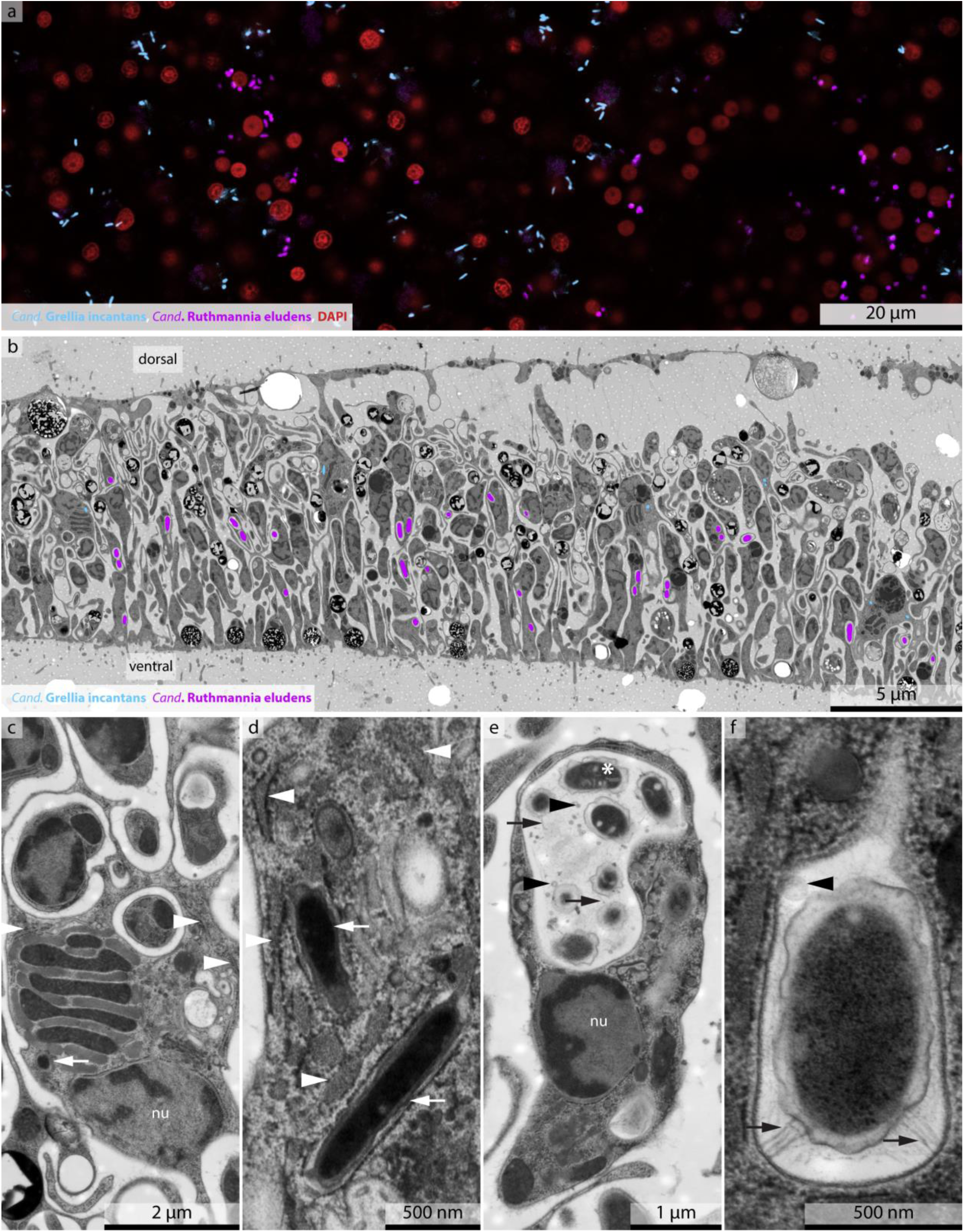
*Cand*. Ruthmannia eludens and *Cand.* Grellia incantans are spatially segregated and specific to two host cell types. **a**, FISH image using probes specific for G. incantans (light blue) and R. eludens (purple); host nuclei are stained with DAPI (red). **b**, TEM image of a cross-section of *Trichoplax* H2 with G. incantans (light blue) and R. eludens (purple) indicated in false color (for raw image data see Supplementary Figure 4). **c** and **d**, TEM image of fiber cells. G. incantans is indicated with white arrows and the rough ER is indicated with white arrowheads. **e** and **f**, TEM image of ventral epithelial cells containing R. eludens. OMVs are indicated with black arrowheads, fimbriae-like structures are indicated with black arrows and internal structures by a white star.

Our correlative FISH and TEM analyses of five *Trichoplax* H2 individuals revealed that the two bacterial symbionts were always intracellular, spatially segregated, and specific to one of the six host cell types (Fig. 3b and Supplementary Figs. 5, 6 and 7). G. eludens was only observed in fiber cells, and was the only bacterium located in these cells (Fig. 3b and Supplementary Figs. 5 and 6). All G. incantans cells were surrounded by a host membrane densely covered with ribosomes (Figs. 3c, 3d and Supplementary Fig. 6) (n=49 symbiont cells in 9 specimens). Similar host structures surrounding the bacteria in other *Trichoplax* lineages were interpreted to indicate that the bacteria reside inside the host’s rER^5^. An alternative interpretation for such host membrane structures was shown for the human intracellular pathogens *Brucella* and *Legionella*, as well as the amoebal midichloriacean parasite *Cand*. Jidaibacter. These bacteria remodel the phagosome surfaces of their hosts to become covered by host ribosomes as an effective strategy for avoiding digestion by their hosts^15,35,36^.

To resolve the sub-cellular architecture of the G. incantans symbiosis, we used high-resolution 3-D TEM tomography to determine if the structures surrounding the symbiont cells were remodeled phagosomes or rER. Our 3-D electron tomographic reconstructions revealed that the ribosome-covered membranes, in which G. incantans occurred, formed networks that were connected to the nuclear envelope, indicating that the structure in which G. incantans is embedded is in fact rER. G. incantans were only observed in the rER, some even within the same rER lumen, and never in other host structures (Fig. 4; Supplementary Fig. 8; Supplementary Video 1). These analyses thus suggest that G. incantans persistently resides in the rER of its host (Fig. 4).

**Figure 4.**
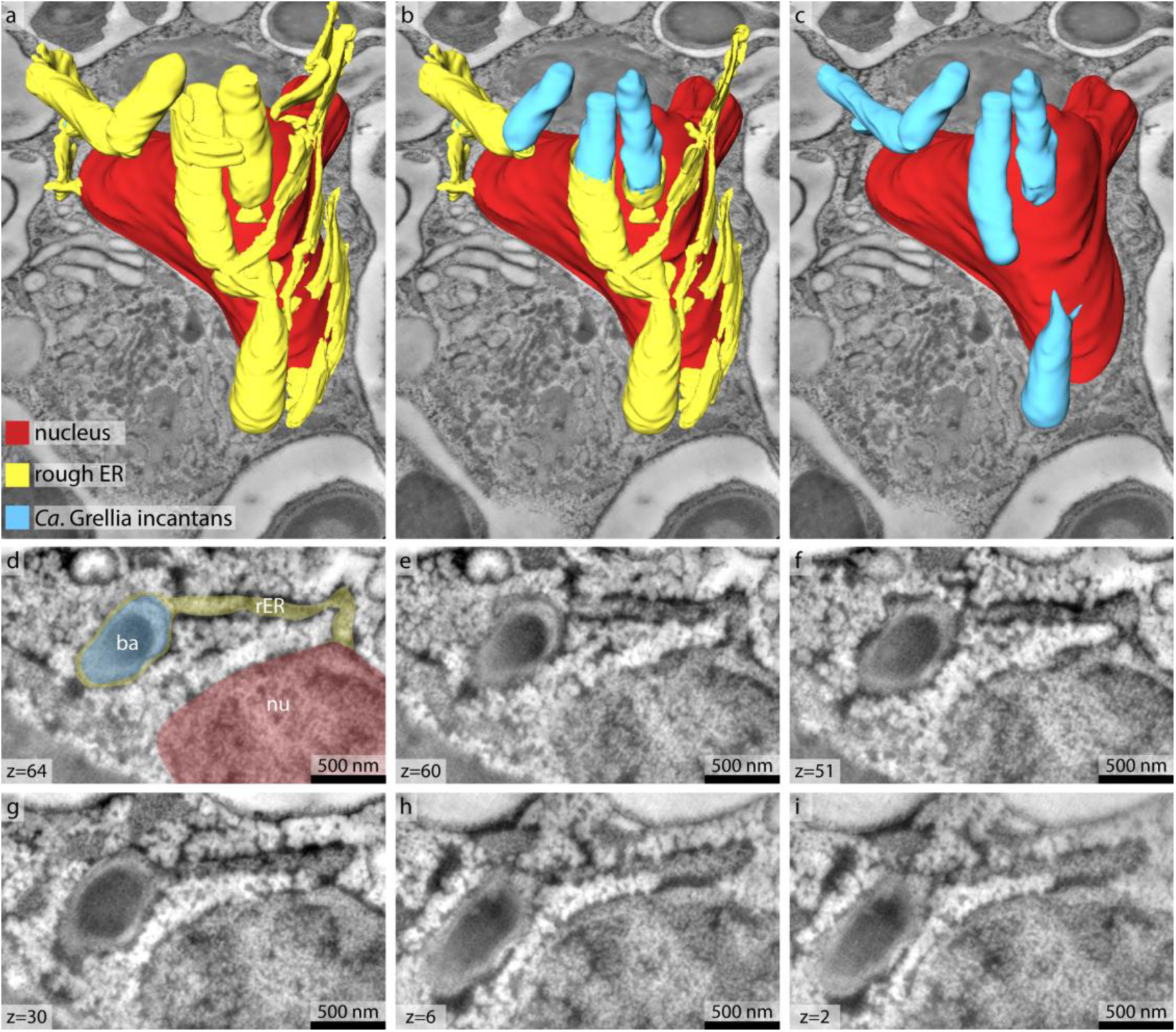
*Ca*. Grellia incantans lives in the rER of *Trichoplax* H2. **a-c**, 3-D volume rendering of reconstructed G. incantans (light blue), rER (yellow) and the nucleus (red) of a fiber cell, superimposed on a virtual slice of the 3D TEM tomography stack. From left to right the rER was virtually removed partially (middle panel) and fully (right panel) to show the symbionts within the rER lumen. No scale bar shown as scale varies with perspective. **d-i**, Selected tomography slices upon which the 3D reconstruction is based show the connection between nucleus, rER and bacteria. For ease of interpretation **d** was false colored with the same color key as in **a**. For raw data see Supplementary Fig. 8.

The second symbiont, R. eludens, only colonized the ventral epithelial cells. These symbionts had a conspicuous, undulated outer membrane, and were always found within cytoplasmic vacuoles of the host, with as many as 15 symbionts co-occurring within a single host vacuole on a single cross-section (Figs. 3e, 3f and Supplementary Fig. 7). These host vacuoles contained numerous membrane-bound vesicles, presumably outer membrane vehicles (OMVs) produced by R. eludens. Conspicuous, thin, electron-translucent, tubular structures appeared to connect the bacterial cells to the host vacuole membrane (Fig. 3f; Supplementary Fig. 7). These fimbriae-like structures may be products of a sec-dependent chaperone-usher (CU) gene set. CU systems are widespread in gram-negative bacteria, and encode essential proteins for the assembly and secretion of adhesive structures^37^. The CU system of R. eludens had remote homologs (25 – 30% amino acid identity) to that of bacteriovorous Deltaproteobacteria (Bdellovibrionales), which use their fimbriae to adhere to their bacterial prey^38^. Both chaperone and usher (PapC and PapD) were expressed, albeit at low levels (Supplementary Table 3).

Bacteria that live inside animal cells are currently known from only six of the 114 recognized bacterial phyla^32^. Despite huge advances in the sequencing of animals from a wide range of phyla and environments that have led to the discovery of numerous lineages of microbiota^29,32^, the number of bacterial phyla with intracellular symbionts has not increased since the characterization of Mycoplasmatales in the early 1960s. Marinamargulisbacteria (UBP8 in Parks *et al.* 2017)^39^ is one of the phylogenetically most remote clades of bacteria, discovered through high-throughput sequencing of environmental samples and advances in binning methods^39^. The remote position of the placozoans in the animal tree of life, together with the technological improvements that enabled the sequencing of individual specimens of these microscopic animals, are likely to have contributed to this late discovery of only the seventh bacterial phylum with intracellular symbionts of animals. The identification of R. eludens from a host that can be easily cultured and investigated using molecular and imaging methods now opens a window to understanding the biology of this enigmatic bacterial phylum.

### *Cand.* Ruthmannia eludens gains nutrition by using lipids degraded by its host (330)

To investigate the physiology of R. eludens, we sequenced the metatranscriptomes of the same single placozoan individuals that were used for metagenomic analyses (n=3), as well as generated metaproteomes from pooled samples of 10 to 30 individuals (n=3). Based on physiological modeling using these expression data, R. eludens is an aerobic chemoorganoheterotroph with a complete TCA cycle that generates energy and biomass from glycerol and the beta-oxidation of fatty acids (Fig. 5a; Supplementary Table 3). The source of the glycerol and fatty acids are most likely lipids derived from the algal diet of the host. Our analyses of the host’s transcriptome revealed that *Trichoplax* H2 expressed several lipases, most likely for the digestion of the algae it feeds on (Supplementary Table 4). These host lipases hydrolyze lipids to glycerol and fatty acids. The genome of R. eludens also encoded lipases. These would allow R. eludens to digest lipids independently of their host. Interestingly, these lipases did not appear to be expressed (Supplementary Table 3).

**Figure 5.**
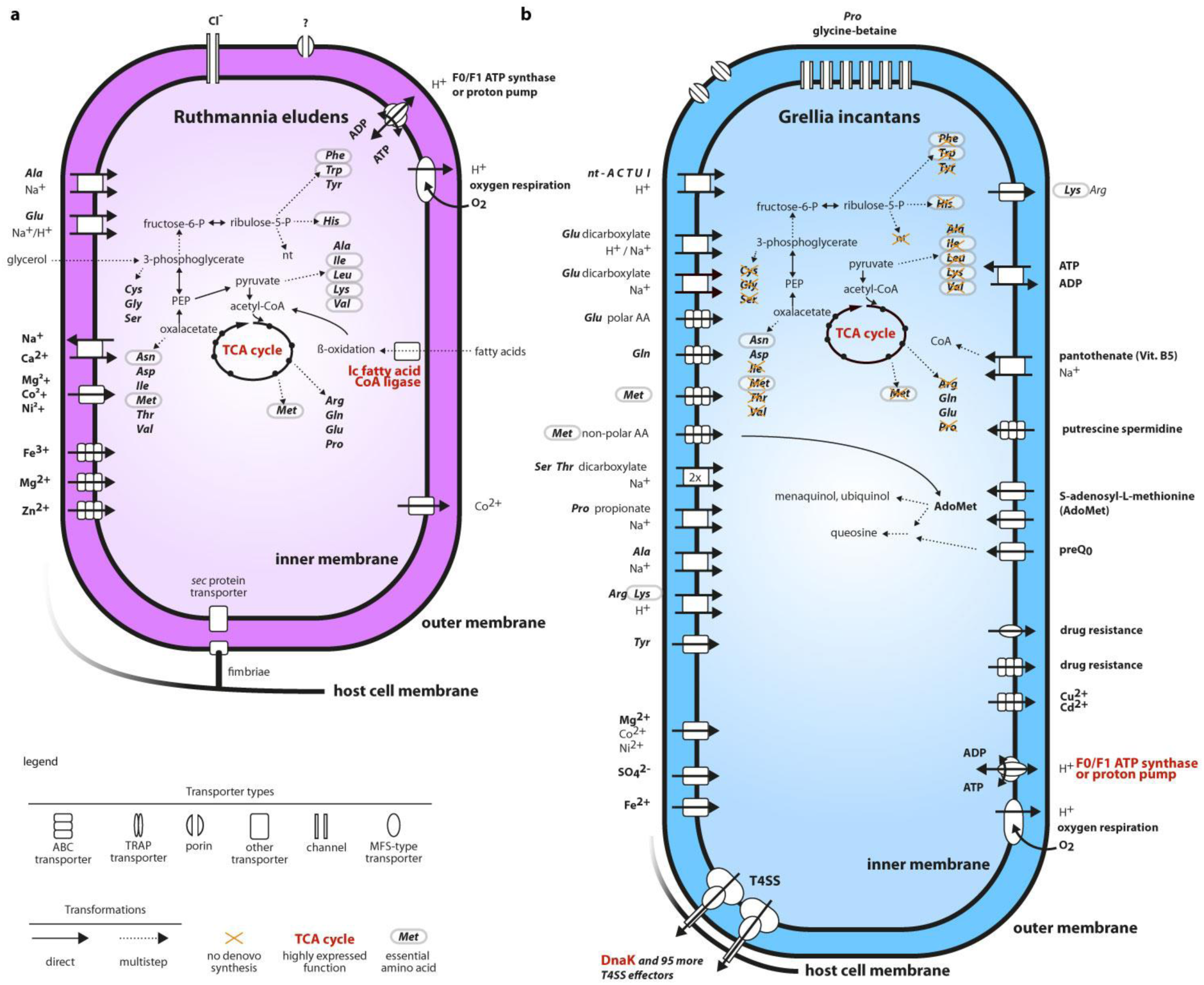
Ruthmannia eludens has versatile biosynthesis pathways, while Grellia incantans depends on the import of most nutrients from its host. Physiological reconstructions based on RAST annotations and Pathwaytools metabolic modelling. Functions that are discussed in the text and highly expressed are indicated in red. **a**, Ruthmannia eludens. **b**, Grellia incantans.

The transfer of glycerol and even-chain fatty acids from the host to R. eludens most likely occurs passively, as they can easily diffuse through cell membranes. We predict that the fatty acids are taken up and activated by R. eludens based on its high expression of a long-chain-fatty-acid-CoA ligase (among the top 25% expressed genes; Fig. 5a; Supplementary Table 3). The fatty acids are then most likely catabolized to acetyl-CoA and respired, as indicated by the expression of all the genes needed for beta-oxidation and the oxidative TCA cycle. The anabolic incorporation of fatty acids is, however, unlikely, as we could not detect the genes for the glyoxylate shunt.

R. eludens encoded genes for synthetizing all nucleotides and amino acids, including the nine amino acids considered essential for animals. However, we found no genomic or transcriptomic indications that R. eludens exports nutrients to its host, for example via amino acid exporters (see Fig. 5a and Supplementary Note 3 for details). Moreover, in our TEM analyses, we found no evidence for the intracellular, lysosomal digestion of R. eludens, such as lamellar bodies or tertiary lysosomes commonly observed in other nutritional symbioses^40,41^. Our ultrastructural analyses did, however, reveal large numbers of putative OMVs in the host vacuole surrounding R. eludens (Figs. 3e, 3f and Supplementary Fig. 8). It is tempting to speculate that the host takes up OMVs produced by R. eludens via phagocytosis and thus supplements its diet, since the host lacks synthesis pathways for essential amino acids. However, the beneficial effects of such putative amino acid provisioning by R. eludens are not clear, given that the animal’s algal diet may contain sufficient amounts of essential amino acids.

### Grellia incantans has the genes for energy parasitism but does not express them

G. incantans, the symbiont that lives in the rER of fiber cells, appears to be a typical Rickettsiales based on genomic features alone, namely a heterotroph that relies on its host for biomass and energy generation (Fig. 5b). The G. incantans genome encoded the hallmark feature for intracellular energy parasites that is present in all Rickettsiales genomes, a fully functional ADP/ATP-translocase for importing ATP from its host^42^. Remarkably, in contrast to all other known energy parasites, we found no evidence for the expression of the ADP/ATP-translocase in G. incantans (Supplementary Table 5). Instead, G. incantans generated ATP with an ATP synthase, and the subunits a and b were highly expressed in the bacterium’s proteome (Supplementary Table 6). Compared to the typical energy-parasitic lifestyle of cytosolic Rickettsiales that rely on ATP imported from their hosts, the ability of G. incantans to synthesize ATP by itself is likely to considerably lower its detrimental impact on its host^43^.

High expression of key genes of the oxidative TCA cycle and the presence of a complete electron transport chain in the genome, with some of the subunits of the electron transport chain among the most highly expressed genes, suggests that the proton gradient for ATP synthesis is fueled by oxidative phosphorylation (Fig. 5b and Supplementary Table 5). An incomplete glycolysis pathway and several importers for α-ketoacids and C4-dicarboxylates suggest that the metabolites respired in the TCA cycle are imported from the host (Fig. 5b).

The genome of G. incantans encoded only a subset of the genes for the *de novo* synthesis of nucleotides. Genes of this subset of the nucleoside/nucleotide biosynthesis as well as genes for parts of the nucleotide conversion pathways were detected in the transcriptome. Similarly, only a subset of the genes for amino acid synthesis were found, none of which were expressed (Fig. 5b; Supplementary Note 4 for details). The apparent lack of amino acid synthesis pathways could be compensated for by a set of 18 importers for amino acids, many of which were expressed. Furthermore, we detected several importers for nucleotides, phosphorus and trace elements in the genome (Fig. 5b). While G. incantans apparently relies on its host for amino acids, it may supply its host with riboflavin. G. incantans expressed the genes for the synthesis of riboflavin (vitamin B2), an essential vitamin that cannot be synthetized by most metazoans. Our analyses of transcriptomic data of the *Trichoplax* H2 host, as well as the genome and proteome of the closely related haplotype H1^18,44^, revealed that both appear to lack the known genes for synthetizing riboflavin (Supplementary Fig. 9). Furthermore, our analyses of the *Trichoplax* H2 transcriptome and the H1 proteome^44^ showed that both haplotypes expressed the enzymes for the conversion of riboflavin to flavin adenine dinucleotide via flavin mononucleotide, and are therefore likely to rely on an external source of riboflavin (Supplementary Table 4). This suggests that by synthetizing riboflavin, G. incantans may supplement the nutrition of its host.

Rickettsiales are known to manipulate their hosts’ cellular biology and evade recognition by its immune system^45^. These manipulations often rely on secretion systems and their secreted effectors. G. incantans encoded two variants of the type IV secretion system (T4SS). The T4SSs are versatile export systems that secrete proteins with a specific C-terminal peptide signature^46^. We detected 96 proteins with T4SS specific C-terminal peptide signatures in the genome of R. incantans, several of which were among the most highly expressed genes. However, many had little homology to well characterized proteins and could therefore not be properly annotated. The three genes with the highest average expression and a T4SS export-peptide signature that could be annotated may be involved in preventing apoptosis. Apoptosis is one of the most common responses of eukaryote cells to bacterial infection^47^, and many pathogenic intracellular bacteria inhibit apoptosis by injecting effector proteins into their hosts through secretion systems. The three annotated genes were LSU ribosomal protein L7/L12, SSU ribosomal protein S11p and the chaperone protein DnaK. While L7/L12 and S11p could not be detected in all three transcriptomes, the highest consistently expressed and annotated protein with a T4SS signature was DnaK, the bacterial homologue to heat shock protein 70 in eukaryotes (Hsp70/Hsp72). Eukaryotic Hsp70 prevents initiation of apoptosis in eukaryotic cells by blocking caspase-9 recruitment to the Apaf-1 apoptosome^47,48^. Eukaryotic Hsp72 has been shown to dampen the unfolded protein response of the rER, a cellular rescue mechanism that is tightly linked to the detection of viral or bacterial interference with eukaryotic protein expression^49^. G. incantans may export DnaK to exploit these two mechanisms and downregulate an immune response of the host. A similar use of DnaK was reported for the alphaproteobacterial pathogen *Brucella*^50^.

G. incantans does not appear to be detrimental, despite the fact that it has to import most of the compounds it needs for generating energy and biomass from its host. Our metagenomic, FISH and TEM data revealed low numbers of symbiont cells in the fiber cells. We estimated the number of G. incantans cells per host cell using metagenomic coverages as proxies of cell abundances for the symbionts and *Trichoplax*. We related the ratio between the host and the symbiont metagenomic abundances to the estimated number of cells in a *Trichoplax* individual as determined previously^14^ (Supplementary Note 5). We estimated that single fiber cells had between 2 – 20 symbionts, numbers that are supported by our FISH and TEM analyses. The total number of G. incantans cells per host individual is thus roughly the same as the number of eukaryotic cells, indicating closely regulated control of bacterial growth by the symbiont, the host or both partners. Pathogen abundances are typically orders of magnitude higher per host cell, often result in rapid exploitation and destruction of host cells and commonly impair host reproduction^51^. The relatively low abundance of G. incantans in *Trichoplax* H2, together with rapid doubling rates of these hosts in our aquaria of 2-3 days, are in stark contrast to virulent pathogenic infections. Moreover, G. incantans appears to generate its own ATP in contrast to all other known energy parasites and modulate its host immune response to prevent apoptosis. It may also supplement its host diet with riboflavin, a potentially beneficial trait when riboflavin availability is limiting for the host.

### Bacterial phylotypes highly similar or identical to G. incantans occur worldwide in aquatic environments

To assess how widespread the two *Trichoplax* symbionts are in other environments and hosts, we surveyed the ∼300,000 publically available amplicon-based 16S rRNA sequence libraries using the IMNGS pipeline^52^. We did not find any sequences related to R. eludens, using a cut-off of 99% identity. In stark contrast, highly similar to identical G. incantans sequences were present in aquatic environments, both marine and limnic, from across the globe (Table 1). Of the 8,026 libraries from aquatic environments, we found sequences that were at least 99% identical to G. incantans in almost ten percent of these libraries (n=845). Out of these 3002 sequences, 1057 sequences were considered identical to the G. incantans sequence and 99.8% of these sequences were attributed to the genus Grellia based on evolutionary placement analysis (Supplementary Fig. 10). This is remarkable for Midichloriaceae, because all other genera were much rarer and present in only 0 – 55 libraries, depending on the genus (Supplementary Table 7). The presence of Grellia phylotypes in such a wide range of environments, including limnic ones, indicates that these bacteria have host ranges beyond placozoans. Indeed, our phylogenetic 16S rRNA analyses showed that sequences that group with the genus Grellia have been found in marine protists (*Eutreptiella*), sea cucumbers (*Apostichopus*), and oysters (*Crassostrea*), as well as in the limnic cnidarian *Hydra oligactis* (see Fig. 1). The *Hydra* sequences came from specimens collected freshly from their natural environments and in animals reared in the laboratory for more than 30 years, indicating the stability of this association in these hosts^53,54^.

The recent realization that human pathogens such as Chlamydiae, Legionellales, and Rickettsiales have closely related relatives that live in hosts ranging from protists to fish from aquatic and soil habitats, has led to a paradigm shift in our view of the ecology and evolution of intracellular bacteria^24,55,56^. G. incantans extends our conceptual understanding of the pervasiveness of such bacteria and shows that a single ‘environmental’ rickettsial genus occurs worldwide in marine and limnic habitats. This remarkable distribution raises the question if all Grellia are host-associated. If G. incantans had a free-living stage, this would be in contrast to all other known Rickettsiales that infect animals ^57^.

## Conclusions

Unlike other animals at the base of the animal tree, such as sponges, cnidarians or ctenophores, Placozoa is the only phylum in which intracellular bacteria have been observed in all individuals and haplotypes investigated. Intracellular symbiosis thus appears to be an invariant trait across this phylum. Our study identifies these bacteria in *Trichoplax* H2, shows that they are found in every specimen examined, and defines the specificity and fidelity to the host cell type in which the symbionts reside.

Although intracellular symbionts are a shared characteristic of all placozoans investigated to date, only little is currently known about the diversity of these symbionts across the 19 cryptic species within this phylum. Our study provides the first insights into how these symbioses may have evolved in two very closely related *Trichoplax* haplotypes, H1 and H2. These two haplotypes putatively separated only decades ago^18^. Intriguingly, their symbioses appear to have followed very different trajectories. While all *Trichoplax* H2 specimens we investigated in this study had the symbiont R. incantans of the Margulisbacteria, neither this symbiont nor any of its close relatives, appears to be present in *Trichoplax* H1 (see Methods). These findings suggest that either: i) the last common ancestor of *Trichoplax* H1 and H2 had a margulisbacterial symbiont that was lost in the H1 lineage; or ii) the last common ancestor of these two host haplotypes did not have a margulisbacterial symbiont, and the H2 lineage acquired this symbiont recently, after separating from H1.

Similarly, the rickettsial symbionts of *Trichoplax* H1 and H2 may have also been acquired independently. The rickettsial symbionts of these two host lineages belong to two different bacterial genera and their 16S rRNA sequences differ by 1.7%. Rates of 16S rRNA divergence in bacteria are estimated to range between 2-11% per 100 million years^58^. Even if these estimates are off by one or even two orders of magnitude, the H1 and H2 symbionts are likely to have diverged from each other at least one million years ago. Assuming that the H1 and H2 hosts separated only decades ago, the vast difference in the time the hosts diverged compared to the divergence time of their symbionts implies that co-speciation could not have occurred. Instead, we envision the following scenarios: 1) The last common ancestor of H1 and H2 had a Grellia-related symbiont. In H1, the Grellia symbiont was replaced by a bacterium from the RETA1 clade, while H2 retained its Grellia symbiont. Or vice-versa, H1 retained its symbiont, and H2 acquired a symbiont from the Grellia lineage. 2) The last common ancestor of H1 and H2 had a symbiont unrelated to the H1 and H2 symbionts. H1 then acquired a symbiont from the RETA1 lineage, while H2 acquired its symbiont from the Grellia genus. Rickettsiales are well-known manipulators of animal sexual reproduction, and it is tempting to speculate that one or multiple infections with Midichloriaceae could have constrained reproductive patterns and possibly shaped the recent divergence between *Trichoplax* H1 and H2^18^. Clearly, future studies of the microbiome of the large number of extant haplotypes are needed to understand more fully the ecology and evolution of symbioses between placozoans and their bacterial symbionts.

## Methods

### Isolation and cultivation

The placozoans were isolated from a coral tank at the Kewalo Marine Laboratory, University of Hawai‘i at Mānoa, Honolulu, Hawai‘i in October 2015 by placing glass slides mounted in cut-open plastic slide holders into the tank for 10 days^11^. Placozoans were identified under a dissection microscope, transferred to 400 ml glass beakers with 34.5 ‰ artificial seawater (ASW) and fed weekly with 2×10^6^ cells ml^−1^ of *Isochrysis galbana* from a log-phase culture. At 25°C in 34.5 ‰ ASW and with a 16:8 hour light/dark regime, doubling times were 2-3 days.

### Nucleic acids extractions

DNA was extracted from two single individuals of the *Trichoplax* H2 cultures using the DNeasy Blood & Tissue Kit (Qiagen) and DNA and RNA from three additional single individuals were extracted using the AllPrep DNA/RNA Micro Kit (Qiagen), according to manufacturer’s protocols with both kits except for the following modifications. Proteinase K digests were performed over night. Elution volumes were halved and all samples were eluted twice, reusing the first eluate. Elutions were carried out with a 10 minutes waiting step before centrifugation.

### DNA and RNA sequencing

Illumina-library preparation and sequencing was performed by the Max Planck Genome Centre, Cologne, Germany. In brief, DNA/RNA quality was assessed with the Agilent 2100 Bioanalyzer (Agilent) and the genomic DNA was fragmented to an average fragment size of 500 bp. For the DNA samples, the concentration was increased (MinElute PCR purification kit; Qiagen) and an Illumina-compatible library was prepared using the Ovation^®^ Ultralow Library Systems kit (NuGEN) according the manufacturer’s protocol. For the RNA samples, the Ovation RNA-seq System V2 (NuGen) was used to synthesize cDNA, and sequencing libraries were then generated with the DNA-library prep kit for Illumina (BioLABS). All libraries were size selected by agarose gel electrophoresis, and the recovered fragments quality assessed and quantified by fluorometry. Per DNA library 14 – 22 million 150 bp paired-end reads were sequenced on a HiSeq 4000 (Illumina), and, for the RNA libraries, 150 bp single-end reads were sequenced to a depth of 42 – 44 million.

### Genome analyses

Full length 16S rRNA gene sequences were reconstructed for each metagenomics and metatranscriptomic library using phyloFlash (https://github.com/HRGV/phyloFlash) from raw reads.

For assembly, adapters and low-quality reads were removed with bbduk (https://sourceforge.net/projects/bbmap/) with a minimum quality value of two and a minimum length of 36; single reads were excluded from the analysis. Each library was error corrected using BayesHammer^59^. A combined assembly of all libraries was performed using SPAdes 3.62 ^60^ with standard parameters and kmers 21, 33, 55, 77, 99.

The reads of each library were mapped back to the assembled scaffolds using bbmap (https://sourceforge.net/projects/bbmap/) with the option fast=t. Scaffolds were binned based on the mapped read data using MetaBAT ^61^. The binning was refined using Bandage ^62^ by collecting all contigs linked to the contig that contained the full-length 16S rRNA gene of the target organism. The bin quality metrics were computed with QUAST ^63^ and the completeness for all bins was estimated using checkM version 1.07 ^64^.

Annotation of the symbiont draft genomes was performed using RAST ^65^ and verified with PSI-BLAST ^66^ for selected genes discussed. Average nucleotide and amino acids identities between genomes ^28^ were calculated with the ANI/AAI matrix calculator (http://enve-omics.ce.gatech.edu/g-matrix). Comparative analyses were conducted using the PATRIC database and services ^67^. Pathway Tools ^68^ in combination with the BioCyc database ^69^ was used to analyse the metabolic capacities of G. incantans and R. eludens. The genomes were screened for secretion systems and effectors using EffectiveDB ^70^.

### Transcriptomic analyses

Adapters and rRNA gene reads were removed from the RNASeq reads using bbduk. Gene expression for each symbiont genome bin and of the host based on the published predicted proteome of *Trichoplax* adhaerens H1 was calculated from RNASeq libraries using kallisto ^71^. Transcription levels were mapped onto metabolic pathways using Pathwaytools ^68^.

### Proteomic analyses

Peptide samples for proteomics were prepared and quantified from two samples of 10 *Trichoplax* each and one sample of 30 *Trichoplax* specimens as described by Kleiner *et al.*^72^ according to the filter-aided sample preparation (FASP) protocol described by Wisniewski *et al.*^73^. In addition to minor modifications as described in Hamann and co-authors^74^, we did not clear the lysate by centrifugation after boiling the sample in lysis buffer. Instead, since the sample size was extremely limited (10 *Trichoplax* specimens = 0.2 ul), we loaded the whole lysate on to the filter units used for the FASP procedure. Centrifugation times before column washes with 100 μl UA were halved as compared to Hamann *et al.*^74^. Peptides were not desalted. Peptide concentrations were determined with the Pierce Micro BCA assay (Thermo Fisher Scientific) following the manufacturer’s instructions.

All samples were analyzed by 1D-LC-MS/MS as described in Kleiner *et al.*^72^ with the modification that a 75 cm analytical column was used. Briefly, the sample containing 30 specimens was measured in technical replicate, for the others the whole sample was used in one analysis. 0.8-3 μg peptide were loaded with an UltiMate^TM^ 3000 RSLCnano Liquid Chromatograph (Thermo Fisher Scientific) in loading solvent A (2% acetonitrile, 0.05% trifluoroacetic acid) onto a 5 mm, 300 µm ID C18 Acclaim^®^ PepMap100 pre-column (Thermo Fisher Scientific). Elution and separation of peptides on the analytical column (75 cm × 75 µm analytical EASY-Spray column packed with PepMap RSLC C18, 2 µm material, Thermo Fisher Scientific; heated to 60 °C) was achieved at a flow rate of 225 nl min^−1^ using a 460 min gradient going from 98% buffer A (0.1% formic acid) to 31% buffer B (0.1% formic acid, 80% acetonitrile) in 363 min, then to 50% B in 70 min, to 99% B in 1 min and ending with 99% B. The analytical column was connected to a Q Exactive Plus hybrid quadrupole-Orbitrap mass spectrometer (Thermo Fisher Scientific) via an Easy-Spray source. Eluting peptides were ionized via electrospray ionization (ESI). Carryover was reduced by to wash runs (injection of 20 µl acetonitrile, 99% eluent B) between samples. Data acquisition in the Q Exactive Plus was done as in Petersen *et al.*^26^.

A database containing protein sequences from the *Trichoplax* host as well as from the two symbionts was used. Sequences of common laboratory contaminants were included by appending the cRAP protein sequence database (http://www.thegpm.org/crap/). The final database contained 13,801 protein sequences. Searches of the MS/MS spectra against this database were performed with the Sequest HT node in Proteome Discoverer version 2.2.0.388 (Thermo Fisher Scientific) as in Petersen *et al.*^26^. For protein quantification, normalized spectral abundance factors (NSAFs)^75^ were calculated per species and multiplied by 100, to give the relative protein abundance in %.

### Phylogenetic and phylogenomic analyses

A 16S rRNA gene database for G. incantans was constructed using the assembled 16S rRNA gene sequence from each metagenomic library, the 20 best BLAST^76^ hits in nr and all other sequences of described Candidatus taxa in the Midichloriaceae. We added the 5 type strains with the best BLAST hit score (5 species of *Rickettsia*) as an outgroup. We also screened the trace reads from the *Trichoplax* H1 genome project for reads containing Midichloriaceae 16S rRNA gene fragments using BLAST^76^, assembled them in Geneious R9 (http://www.geneious.com)^77^ and added the resulting sequence to the database. A similar search for Margulisbacterial 16S rRNA fragments yielded no hits.

The 16S rRNA gene dataset was aligned using mafft^78^, and the phylogenetic tree was calculated using fasttree^79^ with GTR model for nucleotide substitution. The tree was drawn with Geneious^77^.

For G. incantans, the database of genomes for phylogenetic analysis was compiled from all available genomes from the Midichloriaceae as well as representatives for all genera of the Anaplasmataceae and Rickettsiaceae. We also screened the assembly of the *Trichoplax* H1 genome project for contigs that belong to the Midichloriaceae contamination using BLAST^76^ with the G. incantans genome as implemented in Geneious R9 (http://www.geneious.com)^77^. The identified set of contigs corresponded to the set found by Driscoll *et al*.^19^ and were added to the database. We similarly searched for sequences related to R. eludens in the H1 genome project, but no significant hits were detected.

For genome-based alignments of the amino acids of 43 conserved phylogenetic marker genes, the tree workflow as implemented in CheckM was used^64^. For Ruthmannia, the genome bin data was integrated into a taxonomically selected part of the alignment from Hug *et al.* 2016^29^ that covered all Melainabacteria and Cyanobacteria, the WOR-1 and RBX-1 (Margulisbacteria) as well as 5 short branching Firmicutes as an outgroup. The phylogenetic reconstructions of the concatenated alignments were calculated using fasttree with the WAG model for amino acid substitutions and visualized and analyzed using iTOL^80^.

### Tag sequence data analysis

The 16S rRNA gene sequences from G. incantans as well as representative sequences from all characterized midichloriacean *Candidatus* taxa were used as query sequences to search the global collection of microbial tag sequencing library. The search was carried out using the IMNGS service^52^ with a minimal alignment length of 200 bp and a minimal identity of 99%. Identified amplicon libraries were grouped according to their deposited metadata. For the top 10% libraries with the highest number of for sequences from G. incantans, the habitat type (limnic/marine) and geolocation were manually collected in the deposited metadata and the related publications. The detected 16S rRNA reads were aligned to the Rickettsiales dataset using mafft --addfragments and the evolutionary placements in the tree were performed using raxml^81^.

### Transmission electron microscopy

Live specimens were high-pressure frozen with a HPM 100 (Leica Microsystem) in 3 mm aluminum sample holders, using hexane as filler as needed. The samples were transferred onto frozen acetone containing 1% osmium tetroxide and processed using the super quick freeze-substitution method ^82^. After reaching room temperature, the samples were washed three times with acetone and infiltrated using centrifugation, modified after McDonald^83^ in 2 ml tubes sequentially with 25%, 50%, 75% and 2x 100% Agar Low Viscosity resin (Agar Scientific). For this process, the samples were placed on top of the resin and centrifuged for 30 s with a bench top centrifuge (Heathrow Scientific) at 2,000 g for each step. After the second pure resin step, they were transferred into fresh resin in embedding molds and polymerized at 60 °C for 12 hours.

Ultra-thin (70 nm) sections were cut with an Ultracut UC7 (Leica Microsystem) and mounted on formvar-coated slot grids (Agar Scientific). They were contrasted with 0.5% aqueous uranyl acetate (Science Services) for 20 min and with 2% Reynold’s lead citrate for 6 min before imaging them at 20-30 kV with a Quanta FEG 250 transmission electron microscope (FEI Company) equipped with a STEM detector using the xT microscope control software ver. 6.2.6.3123.

For electron tomography 300 nm serial sections were placed on formvar coated 2×1mm slot grids and stained with uranyl acetate and lead citrate. 30 nm gold fiducials were applied on both sides of the slot grid. Dual-axis tilt series (±60°, step size 1°) were acquired with a FEI Tecnai F30 300kV electron microscope equipped with an Axial Gatan US1000 CCD camera. SerialEM software was used for the automated tomographic tilt series acquisition^84^. Alignment and reconstruction of the tilt series were carried out with IMOD^85^. The serial tomograms were aligned with TrakEM2^86^ in Fiji^87^ and visualization and segmentation were carried out using the software Amira 3D.

### Fluorescence in situ hybridization

We used the arb-silva database 128^88^ and the arb PROBE_DESIGN tool (the arb software package)^89^ to design two FISH probes for each symbiont that were specific to their 16S rRNA sequences (Supplementary Table 2). We confirmed the specificity of the probes by comparing their sequences to all available sequences in the arb-silva 128 database and RDP (Ribosomal Database Project) rel.11.5^90^. The most specific probe to R. eludens had 2 mismatches to first non-target hit sequences, the most specific probe for G. eludens also matches the 6 most closely related Grellia sequences, detailed results are presented in Supplementary Table 2.

Specimens were fixed on coverslips with 2% formaldehyde and 0.1% glutaraldehyde in 1.5X PIPES, HEPES, EGTA and MgCl2 (PHEM) buffer modified from Montanaro *et al.*^91^ at 4°C for 12 hours. After three washing steps in 1.5X PHEM buffer the samples were stored in 70% ethanol until use. Samples were rehydrated in phosphate buffered saline (PBS) and hybridization was performed according to Manz *et al.* ^92^. Mono-labeled-, DOPE-^93^ or MIL-^94^ probes (Supplementary Table 2) at a concentration of 8.4 pmol/µl were diluted with hybridization buffer containing 35% formamide, 900 mM NaCl, 20 mM Tris/HCl and 0.01% SDS at a ratio of 15:1. Whole animals were incubated in 30 µl of the probe/hybridization buffer mix at 46°C in 250 µl PCR tubes for 3-4 hours, followed by a 30 minute washing step in washing buffer containing 700 mM NaCl, 20 mM Tris/HCl, 5 mM EDTA and 0.1% SDS. After a 10 minute washing step in PBS, the animals were stained with DAPI for 30 minutes, washed twice again in PBS and mounted on glass slides in Vectashield mounting medium.

To test the probes designed for this study, 30 clonal individuals of *Trichoplax* H2 were pooled, fixed as described above, homogenized by sonication and applied to a filter. The parts of the filter were then tested with different formamide concentrations and the optimal formamide concentration was determined.

Fluorescence images were taken with a Zeiss LSM 780 equipped with a plan-APROCHROMAT 63X/1.4 oil immersion objective using the ZEN software (black edition, 64bits, version: 14.0.1.201) (Carl Zeiss Microscopy GmbH).

## Data availability

The metagenomic and metatranscriptomic raw reads and assembled symbiont genomes are available in the European Nucleotide Archive under Study Accession Number PRJEB30343

The mass spectrometry metaproteomics data and protein sequence database were deposited in the ProteomeXchange Consortium^95^ via the PRIDE partner repository with the dataset PXD012106

The TEM 3D reconstruction data was deposited in figshare, the aligned tomography slices used for the reconstruction shown in Figure 4 are available at https://figshare.com/s/886b869a9ada0264ffb2 (doi 10.6084/m9.figshare.7429793).

## Acknowledgments

This study was funded by the Max Planck Society with additional support from the Gordon and Betty Moore Foundation Marine Microbial Initiative Investigator Award (Grant GBMF3811) to N.D., grants to M.H. from the Gordon and Betty Moore Foundation (no. 5009) and the U.S. Office of Naval Research grant no. N00014-15-1-2658, by the German Academic Exchange Service DAAD (T.H.) and the NC State Chancellor’s Faculty Excellence Program Cluster on Microbiomes and Complex Microbial Communities (M.K.). We thank M. Strous for access to proteomics equipment and A. Kouris for LC-MS/MS operation. The purchase of the proteomics equipment was supported by a grant of the Canadian Foundation for Innovation to M. Strous.

We thank the Electron Microscopy Facility of the MPI-CBG, the Max Planck-Genome-centre Cologne and the Core Facility Cell Imaging and Ultrastructure Research of the University of Vienna for technical support.

The authors thank C. Peters for nucleic acids extractions, and C. Peters, M. Meyer and W. Ruschmeier for support with the *Trichoplax* cultivation, B. Nedved for the support in the field and G. Bennett, T. Erb, L. Schada von Borzyskowski and P. A. Chakkiath for discussions on symbiont physiology.

## Author contributions

M.H., N.D., M.MF-N. and H.G-V. conceived the study. H.G-V. sampled and cultivated the organisms, performed the assemblies, genome and transcriptome analyses, tag-sequencing analyses and phylogenetic analyses. H.G-V. reconstructed symbiont physiology with the help of M.L. and M.K. M.K, T.H generated proteomic data, and H.G-V. and M.K. analyzed the proteomic data. N.L. performed the fluorescence microscopy, electron microscopy and electron tomography and subsequent data analysis and three-dimensional reconstruction. H.G-V. and N.L. wrote the manuscript with support from N.D., M.MF-N. and M.H. All authors revised the manuscript and approved the final version.

## Supplements

Note: Supplementary Notes 1-5, all Supplementary Figures, Methods and references are provided in a supplementary PDF file.

**Supplementary Note 1 - Description of *Cand*. Grellia incantans**

**Supplementary Note 2 - Description of *Cand*. Ruthmannia eludens**

**Supplementary Note 3 - Ruthmannia eludens physiology**

**Supplementary Note 4 - Grellia incantans physiology**

**Supplementary Note 5 - Metagenomics based symbiont cell number estimates**

**Supplementary Figure 1 – Based on the mitochondrial 16S rRNa the Kewalo *Trichoplax* lineage is a haplotype H2**

**Supplementary Figure 2 – Full length 16S rRNA based diversity of bacteria associated with 5 single individuals of the Kewalo *Trichoplax* H2 lineage**

**Supplementary Figure 3 – Autofluorescence of *Trichoplax* H2**

**Supplementary Figure 4 - Fluorescence in-situ hybridization of the two bacterial phylotypes present in *Trichoplax adhaerens* H2**

**Supplementary Figure 5 – Transmission electron microscopic raw image data used for the false coloration shown in Figure 3d**

**Supplementary Figure 6 – Transmission electron microscopy images of fiber cells and the localization of G. incantans**

**Supplementary Figure 7 – Transmission electron microscopy of ventral epithelial cells and localization of Ruthmannia incantans**

**Supplementary Figure 8 - Electron tomography of Grellia eludens**

**Supplementary Figure 9 - Riboflavin KEGG map of H1 genome and proteome**

**Supplementary Figure 10 – EPA of Grellia matching SRA sequences in midichloriaceae 16S rRNA gene tree**

**Supplementary Table 1 – The microbiome is dominated by Grellia incantans and Ruthmannia eludens**

**Supplementary Table 2 - Overview of the FISH probes used**

Supplementary Tables 3 – 7, Supplementary Video 1 and Supplementary Dataset 1 are provided as separate files

**Supplementary Table 3 – Ruthmannia eludens transcriptome**

**Supplementary Table 4 – *Trichoplax* H2 transcriptome analysis**

**Supplementary Table 5 – Grellia incantans transcriptome**

**Supplementary Table 6 – Grellia incantans proteome**

**Supplementary Table 7 – Tag sequencing libraries with hits from Midichloriaceae.**

**Supplementary Video 1 – Rendering of 3D reconstruction**

**Supplementary Dataset 1 – Aligned tomography stack**

