## Supplementary Information for "The *Trichoplax* microbiome: the simplest animal lives in an intimate symbiosis with two intracellular bacteria"

#### Contents

Contains or links to

Supplementary Notes 1 – 5

Supplementary Methods

Supplementary Figures 1 – 10

Supplementary Video 1

Supplementary Tables 1 – 7

Supplementary Datasets

Supplementary references

#### Supplementary Note 1 - Description of *Cand*. Grellia incantans

We name the *Candidatus* genus Grellia in honor of Karl G. Grell who rediscovered the placozoans after they had been erroneously dismissed as larvae of hydromeduse[1,2]. The species epithet incantans (pres. part. of Latin, *incantare* – bewitching or enchanting) refers to the ability of these bacteria to associate with hosts across the globe and their presence in both limnic and marine environments, possibly using the same book of songs for their wide range of environmental interactions.

*Cand*. Grellia incantans (from here on G. incantans) was co-isolated from a marine aquarium at the Kewalo Marine Laboratories, Honolulu, Hawaii together with its host, *Trichoplax* haplotype H2. It was detected in *Trichoplax* H2 hosts using fluorescence *in situ* hybridization (FISH) with the probes listed in Supplementary Table 2. G. incantans has not been cultivated outside of the *Trichoplax* H2 host. The cells are gram-negative small and thin rods (max. length 1200 nm, avg. length 541±359 nm, max. width 326 nm, avg. width 251±33 nm, n=19 cells). The cells are embedded in the host rough endoplasmic reticulum that is densely lined with ribosomes.

The recovered genome bin that represents the complete bacterial chromosome is 1.26 Mb with an estimated completeness of 98.9% based on 108 conserved bacterial marker genes and no detectable contamination. Compared to the full genome of G. incantans, the 181 coding sequences recovered for the *Trichoplax* H1 phylotype had an average amino acid identity (AAI) of 64.38 % and had an average of 0.16 substitutions per site across 43 highly conserved genes. The AAI value is below the threshold of 65% AAI for genomic similarity within genera. The AAI data and the phylogenomic results support the 16S rRNA-gene based interpretation that the rickettsial sequences detected in *Trichoplax* H1 belong to a separate genus from the G. incantans found in *Trichoplax* H2.

#### Supplementary Note 2 - Description of *Cand*. Ruthmannia eludens

The *Candidatus* genus name Ruthmannia was chosen after August Ruthmann who devoted a significant part of his career to research on placozoans. The *Candidatus* species epithet eludens (pres. part. of Latin, *eludo* – escaping or fooling) refers to how well this symbiont has escaped detection compared to all other intracellular symbionts in metazoans, and how difficult the interpretation of a large part of its genome is, as there are no characterized homologs.

*Cand.* Ruthmannia eludens (from here on called R. eludens) was recovered from a marine aquarium at the Kewalo Marine Laboratories, Honolulu, Hawaii together with its host, *Trichoplax* H2. R. eludens was detected in *Trichoplax* H2 hosts using FISH with the probes listed in Supplementary Table 2. R. eludens has not been cultivated outside of the *Trichoplax* H2 host. The cells are Gram-negative rods (max. length 1158 nm, avg. length 669±233 nm, max. width 469 nm, avg. width 373±46 nm, n=37 cells). The cells are embedded in a host vacuole, and appear to be anchored to the host membrane with fimbriae. We assembled a 1.5 Mb metagenomic bin for R. eludens. Despite the small genome size, the genome draft has a computed completeness of 98.3% based on 108 conserved bacterial marker genes and no detectable contamination.

### Supplementary Note 3 - Ruthmannia eludens physiology

#### Transport across membranes

The only importers that could be annotated were for two non-essential amino acids (alanine and glutamate) and for cations important for metalloproteins ($Ca^{2+}$, $Mg^{2+}$,$Co^{2+}$, $Ni^{2+}$, $Mn^{2+}$, $Zn^{2+}$ and $Fe^{3+}$). The alanine, $Fe^{3+}$, and $Mn^{2+}$ importers were expressed, suggesting that the intracellular R. eludens, to some degree, rely on their host for acquiring nitrogen, iron and manganese.

To interact with the host cell, R. eludens must be able to export signaling molecules. Among the top ten most expressed genes was an outer membrane barrel protein that might serve as a porin, but its substrates could not be resolved, as for many other exporters. The substrate could only be identified for a single exporter gene, the cobalt efflux protein *corC*: Together with the anaerobic cobalt chaperone *cbiX*, *corC* could confer cobalt resistance, but this exporter was not expressed. The expressed SecAYG and YidCD proteins likely form an inner membrane protein assembly system that can translocate proteins from the cytoplasm to the periplasm, but none of the canonical secretion systems to export proteins across the outer membrane and into host cells were present in the genome.

#### Supplementary Note 4 - Grellia incantans physiology

As in the genomes of the tick and amoeba midichloriaceans, the G. incantans genome contained a cbb3 cytochrome oxidase typical for microaerophilic organisms.

##### Flagellar apparatus

A set of 35 genes encoding a fully functional flagellum, including all cytoplasmic, cell envelope-anchored and extracellular structural proteins and the necessary regulatory factors, was present in the genome. We never observed flagella in our electron microscopic analyses of nine host individuals. The presence of transcripts from genes for the cytoplasmic and cell-envelope anchor of the flagellum and a high expression of the filament-protein flagellin indicates that at least a part of the G. incantans population assembles or maintains flagella. This corresponds to the observed expression of flagella in other Rickettsiales during their intracellular stage[3]. A homologous set of flagellar genes was also found in the much larger genome of *Cand*. Jidaibacter, and was interpreted as an ancient feature of this bacterium[4]. In contrast, in *Cand*. Midichloria the set of flagellar genes is incomplete, and the flagellum likely is no longer functional[4]. Three genes of the flagellar apparatus were also detected in the partial rickettsial genome from the *Trichoplax* haplotype H1 genome (RETA1)[5]. As G. incantans and RETA1 belong to the sister clades of *Cand*. Jidaibacter and *Cand*. Midichloria (Figure 1a), it is parsimonious to assume that the last common ancestor (LCA) of the Midichloriaceae had a functional flagellum and was motile. Indeed, ultrastructural imaging and genomic evidence across all families of Rickettsiales suggests that this was the case for the LCA of all Rickettsiales[6].

##### Transport across membranes

G. incantans most likely imports most amino acids, including all that are essential to the *Trichoplax* host, as full amino acid synthesis pathways were only present for aspartate, asparagine, glutamate and glutamine. We identified importers for at least ten amino acids including the essential amino acids methionine and lysine, of which two, for methionine and glutamate, were highly expressed. Typical for intracellular Rickettsia, the genome featured a large array of other importers, e.g. for nucleotides, co-factors like pantothenate (vitamin B5) and S-adenosylmethionine (AdoMet) or inorganics (magnesium, sulfate, iron). In contrast to the versatile importers, the G. incantans genome encoded only five exporters, a copper ABC-transporter, three multidrug exporters and a lysine exporter, but we did not find transcripts for any of these genes. The predicted import of nucleotides and amino acids as well as most other essential metabolites for energy generation and cell homeostasis indicates that G. incantans is dependent on its hosts for biomass formation.

#### Protective polyamines

In the Rickettsiaceae, millimolar concentrations of the polyamines putrescine and spermidine essential for DNA stabilization and protein biosynthesis, have been observed[7]. The expression of the polyamine importer PotABCD in G. incantans indicates that these protective molecules are also important in Midichloriaceae and likely constitute a conserved trait across Rickettsiales.

#### Antioxidants

Compared to *Cand*. Jidaibacter and *Cand*. Midichloria, which appear to have a very limited distribution, one of the factors that contributes to Grellia's wide habitat range could be that it has two variants of the potent antioxidant alkyl hydroperoxide reductase[8]. Similar to the multiple variants in *Bacillus subtilis*, these two variants could have different substrate specificities or expression patterns that would make G. incantans more tolerant against oxidative stress than other Midichloriaceae[9,10].

#### Type IV secretion systems effectors and the mitochondria of fiber cells

The mitochondria in all fiber cells have an enlarged morphology and are alternatingly stacked with vesicles, forming so-called mitochondrial complexes that are a key feature of placozoan fiber cells[2]. G. incantans is specific to the fiber cells and likely present in all of these cells, but we could not find genomic evidence that it is involved in the formation of the aberrant mitochondria, despite the fact that two effectors with mitochondrial signal peptides are highly expressed. The first predicted effector is a conjugative plasmid relaxase associated to the transfer of DNA via the type IV secretion system and unlikely to affect mitochondrial morphology. The second is a small protein (134 amino acids) with no characterized homologues that is conserved across Midichloriaceae. As neither *Cand*. Jidaibacter nor *Cand*. Midichloria induce mitochondrial complexes in their host cells, its function remains to be shown.

#### Supplementary Note 5 - Metagenomics based symbiont cell number estimates

The read coverage in metagenomes is directly related to copy numbers of chromosomes and therefore to cell numbers. The coverage for both symbiont chromosomes in the five metagenomes was variable, but as low or lower than for the host nuclear genome, down to more than 10x lower. An average *Trichoplax adhaerens* individual was estimated to consist of approximately 50.000 host cells [11] and we estimated similar cell numbers for the morphologically indistinguishable H2 haplotype. Using these estimates, we translated the metagenomics coverage ratios to cell counts of 5000-50000 intracellular bacteria for each of the two symbionts. G. incantans resides in the fiber cells that extend throughout the whole animal as a connective cell layer[11] (Fig. 3). Given that on average only 5% of the cells in a host individual are fiber cells[11] this corresponds to 2 - 20 symbiont cells per fiber cell.

### Supplementary Methods

#### Host mitochondrial 16S rRNA gene phylogenetic analyses

The metagenomic assembly (see Methods in main text) was screened for the contig containing the mitochondrial 16S rRNA gene (m16S) using BLAST as implemented in Geneious R11. The gene was extracted from the contig and aligned together with a database of publically available m16S sequences using mafft in G-Insi mode. The phylogenetic tree was reconstructed using fasttree v2.1.5[12] with a GTR model, 20 rate categories and Gamma20 likelihood optimization, generating aLRT values for node support. The tree was drawn with Geneious[13]. The tree was rooted with clade A placozoans[14].

#### Bacterial diversity 16S rRNA gene phylogenetic analyses

For the 16S rRNA gene database of all phylotypes recovered, the phyloFlash (https://github.com/HRGV/phyloFlash) assembled full length SSU genes of all samples were collected. The dataset was aligned and phylogenetic trees were calculated and visualized as for the host m16S dataset above. The tree was rooted with the Eukarya and only the bacterial part of the tree was shown.

#### Supplementary Figures

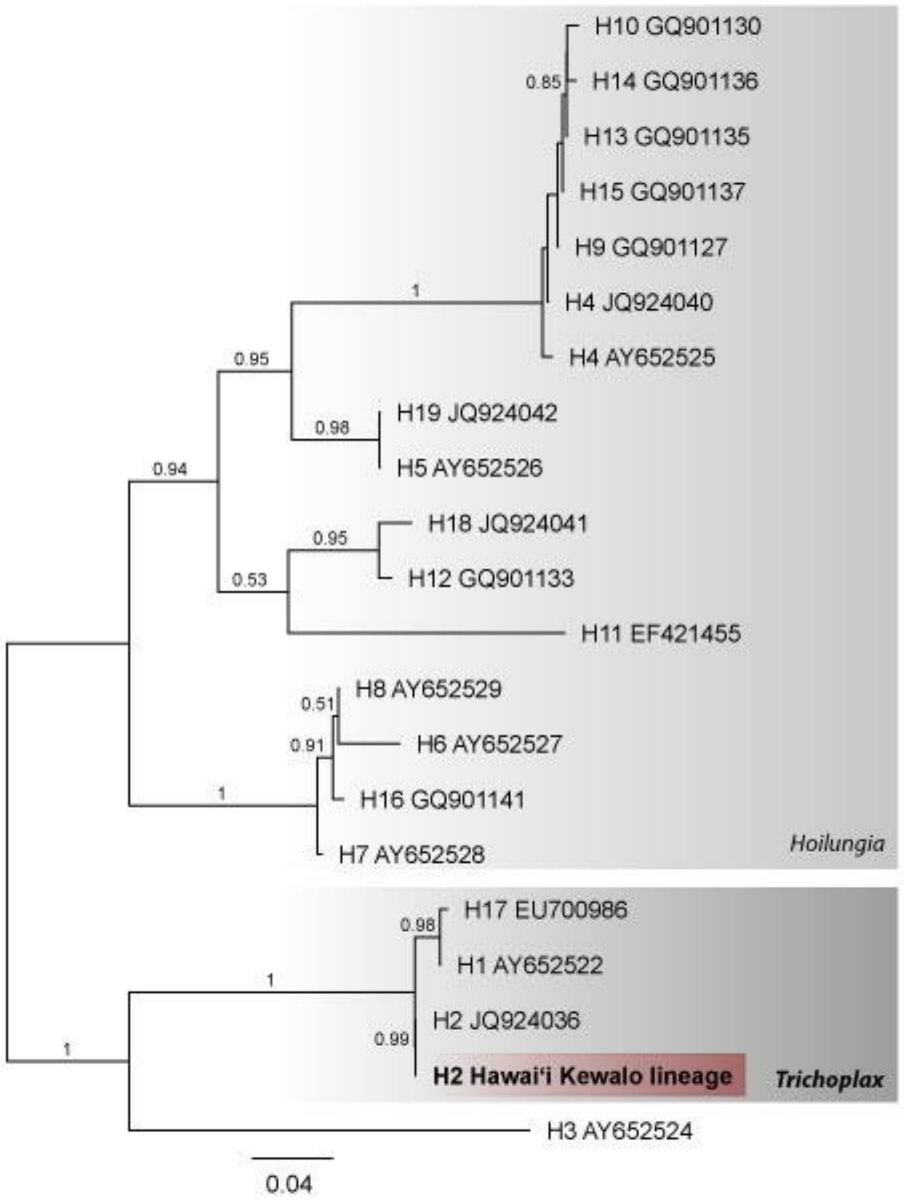


**Supplementary Figure 1 – Based on the mitochondrial 16S rRNA, the Kewalo *Trichoplax* is haplotype H2**

Phylogenetic tree based on the alignment of the mitochondrial large subunit rRNA (16S), support values below 0.5 are not shown. Scale bar indicates substitutions per site. The placozoan genera, haplotypes and the accession numbers are indicated.

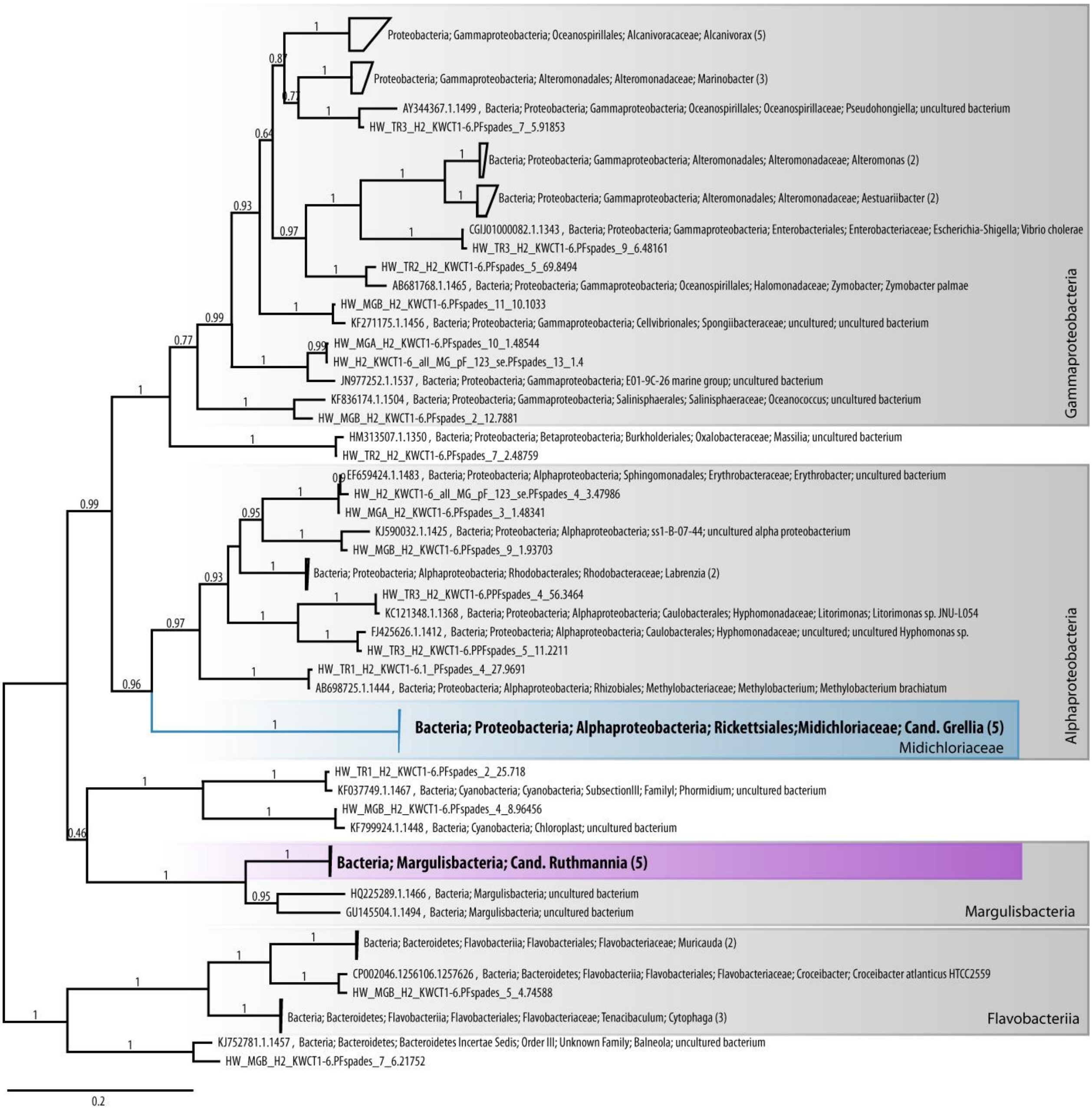


#### Supplementary Figure 2 – Full length 16S rRNA based diversity of bacteria associated with five single individuals of the Kewalo *Trichoplax* H2 lineage

Phylogenetic tree based on the alignment of the bacterial 16S rRNA gene. Scale bar indicates substitutions per site. Major bacterial lineages that were present in at least three samples are indicated in grey.

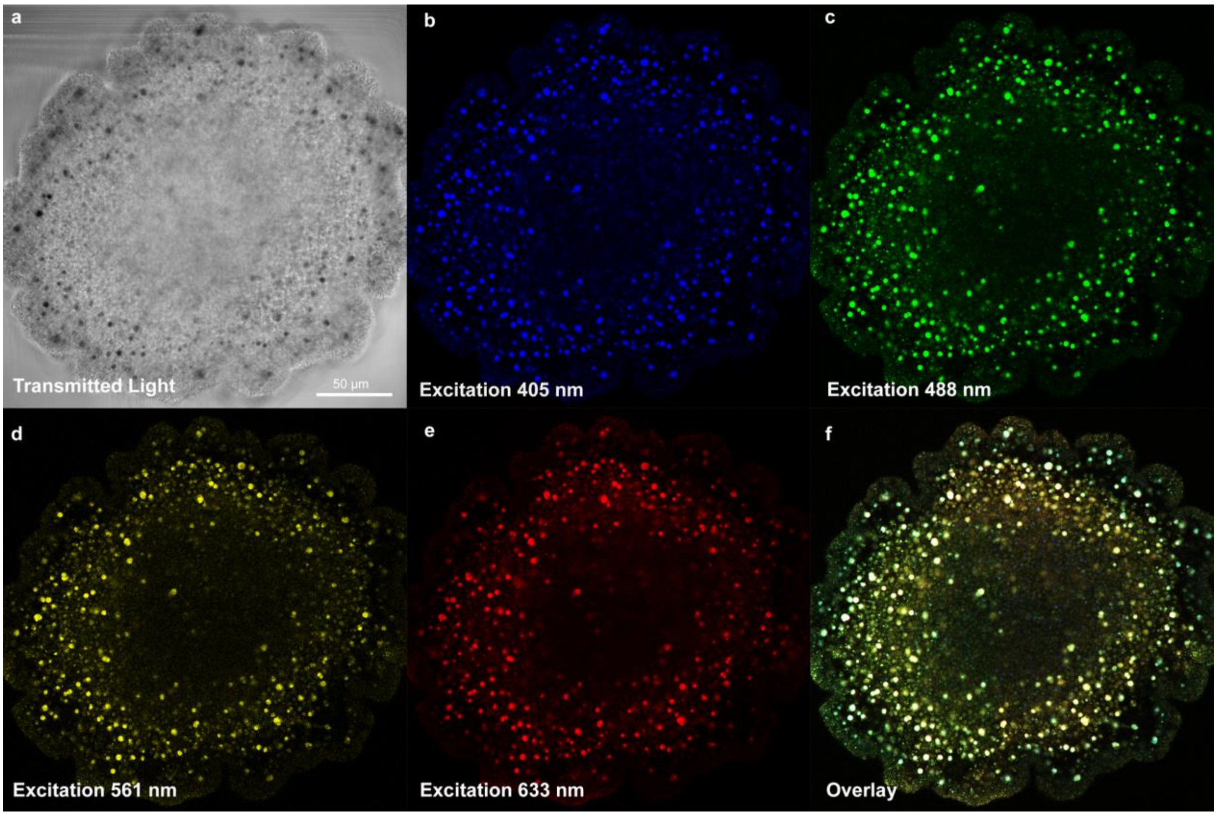


#### Supplementary Figure 3 – Autofluorescence of *Trichoplax* H2

**a**-**f,** Images of the same whole mount *Trichoplax* H2 specimen **a,** Transmitted light image of the animal **b**-**e**, Autofluorescence induced by the most commonly used laser-lines of a confocal laser scanning microscope. **f**, Overlay of **b**-**e**

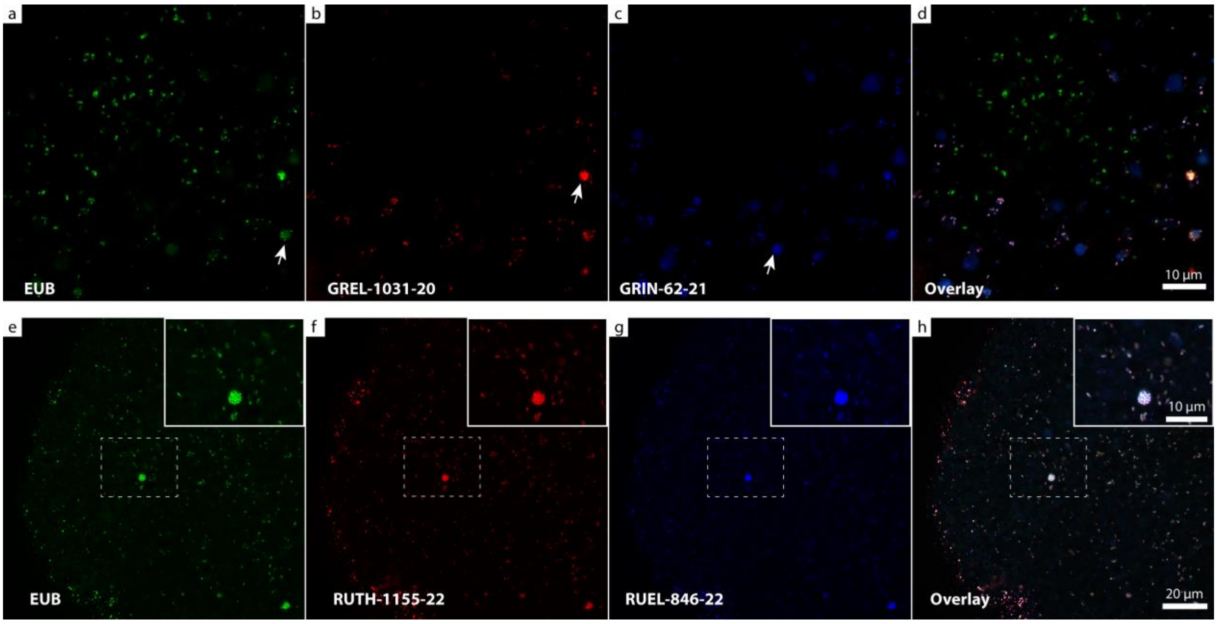


#### Supplementary Figure 4 - Fluorescence in-situ hybridization of the two *Trichoplax* H2 symbionts

**a** and **e,** Eubacterial probe **b** and **c**, Probes specific for G. incantans **f** and **g,** Probes specific for R. eludens **d** and **h,** Overlay of the three individual channels. White arrows in **a**-**c** show areas of high autofluorescence, originating from the food vacuole of the fiber cells. Inset in **e**-**h** shows a higher magnification of the area indicated with a dotted rectangle.

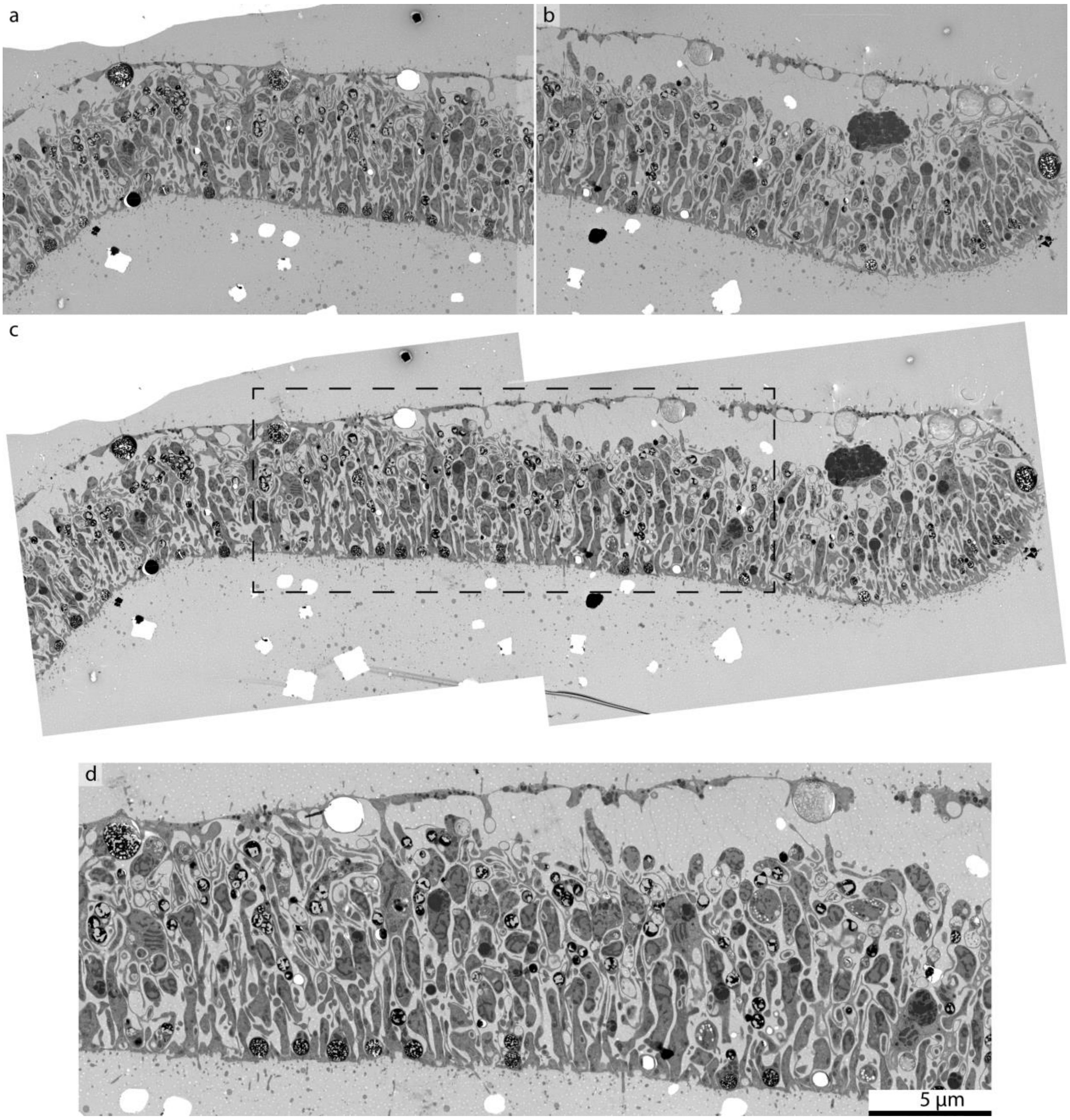


#### Supplementary Figure 5 – Transmission electron microscopic raw image data used for the false coloration shown in Figure 3d

**a** and **b,** The two original images **c,** The combined image **d,** Detail used in the main figure 3d (indicated with a dotted rectangle in **c**).

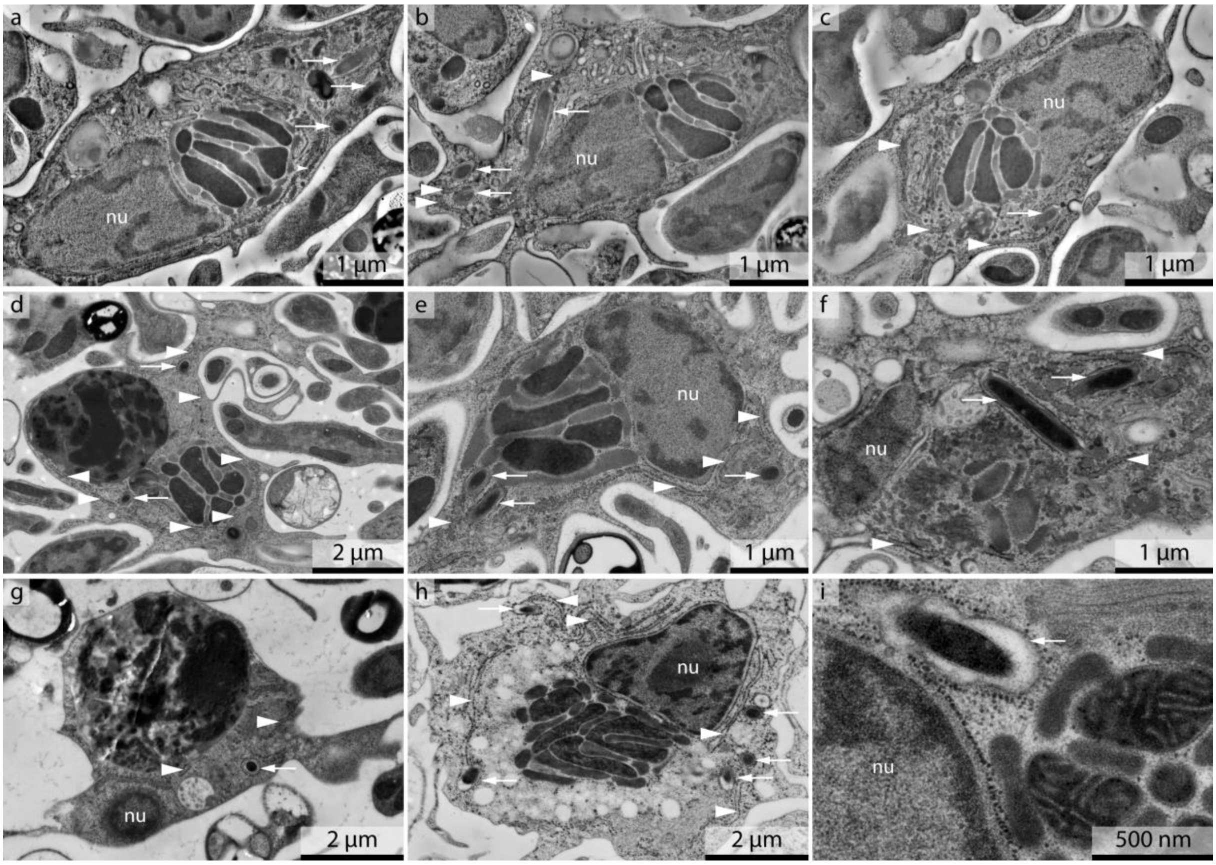


#### Supplementary Figure 6 – Transmission electron microscopy images of fiber cells and the localization of G. incantans.

**a**-**c,** Individual slices extracted from a tomogram **d**-**f,** Thin sections of epon embedded animals **g**-**i,** Thin sections of an LR-White embedded animal. nu indicates the nucleus, arrowheads point towards the rough endoplasmic reticulum and arrows point towards G. incantans located within the rER

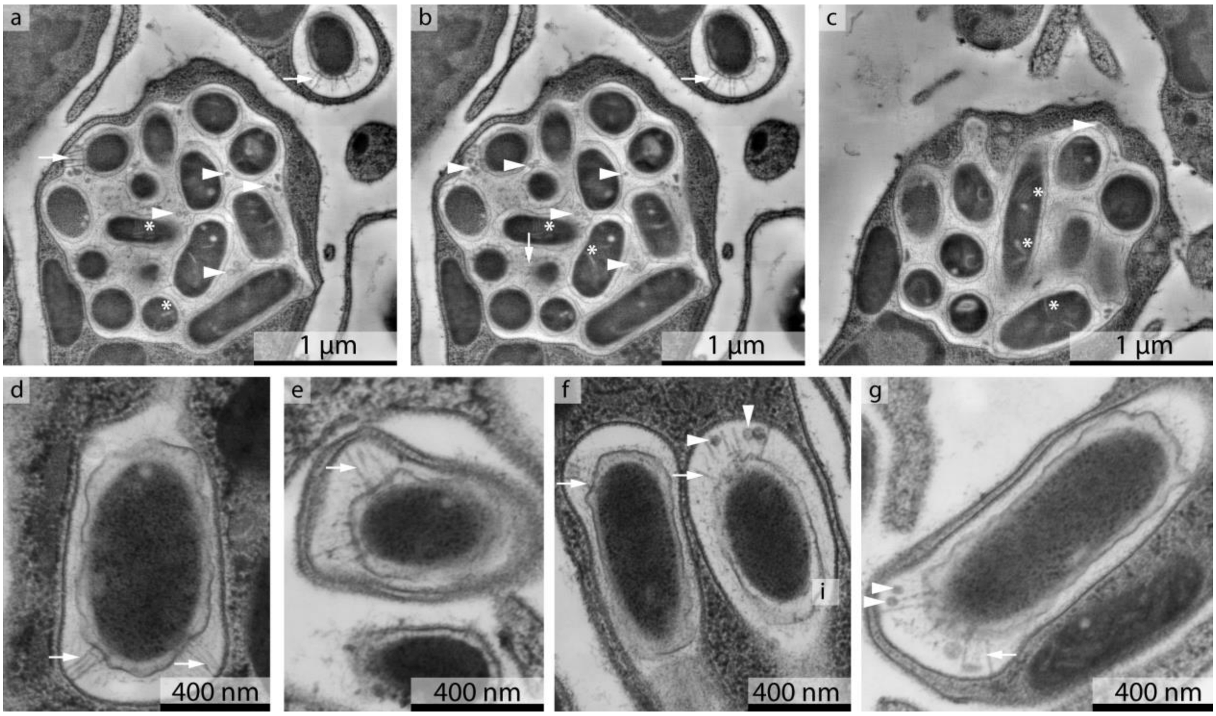


**Supplementary Figure 7 – Transmission electron microscopy of ventral epithelial cells and localization of Ruthmannia eludens**

**a**-**c,** Individual slices extracted from a tomogram showing R. eludens in the ventral epithelial cell **d**-**g,** R. eludens in higher magnification. Arrowheads point to outer membrane vesicles, arrows point towards the fimbriae like structures and the asterisk indicates internal structures in R. eludens.

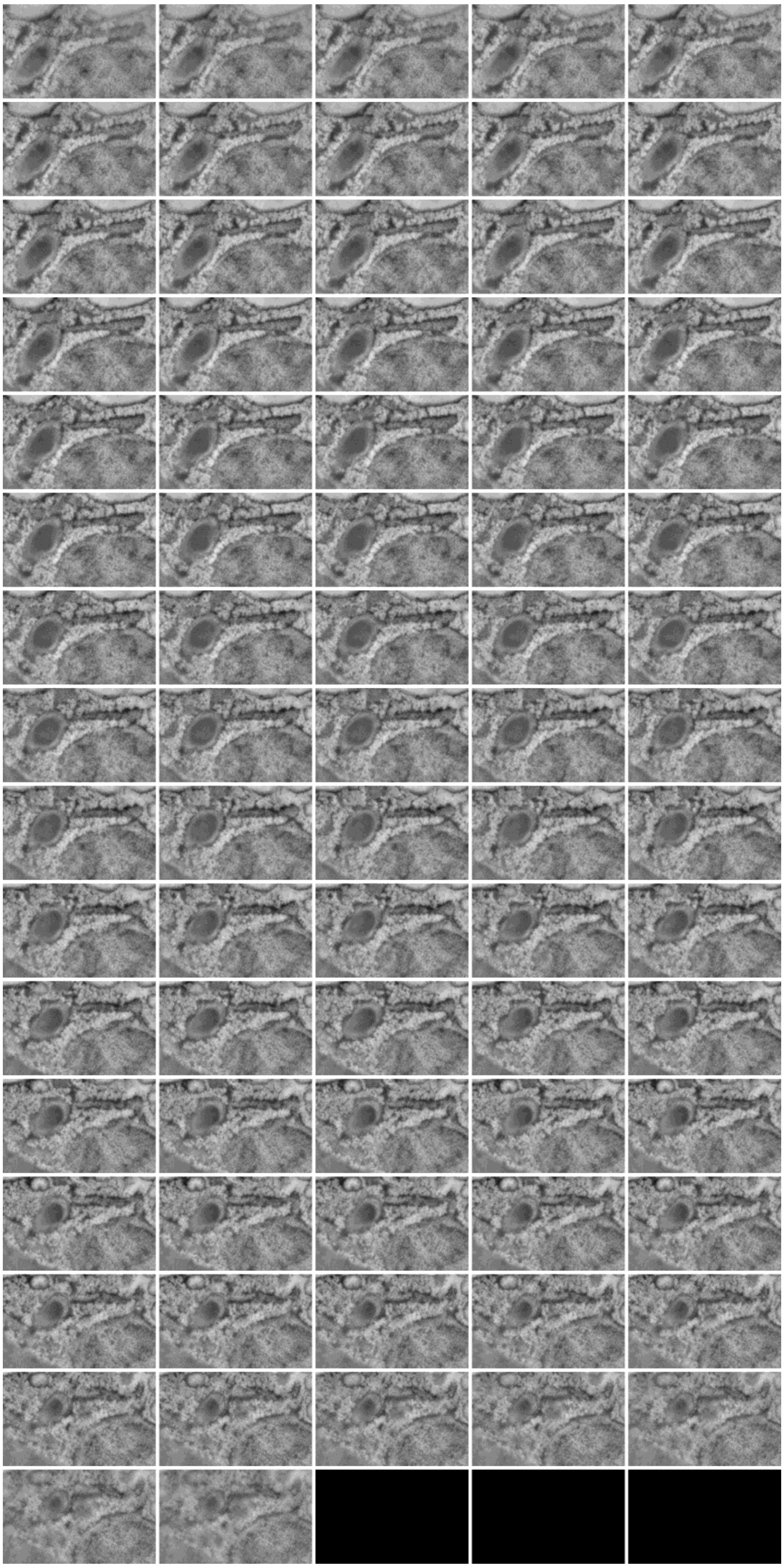

#### Supplementary Figure 8 - Electron tomography of Grellia incantans

Cropped detail from a single tomogram of a 300 nm section, showing G. incantans surrounded by rER, which in turn is connected to the nuclear membrane. Six individual slices of this tomogram were shown in Figure 4.

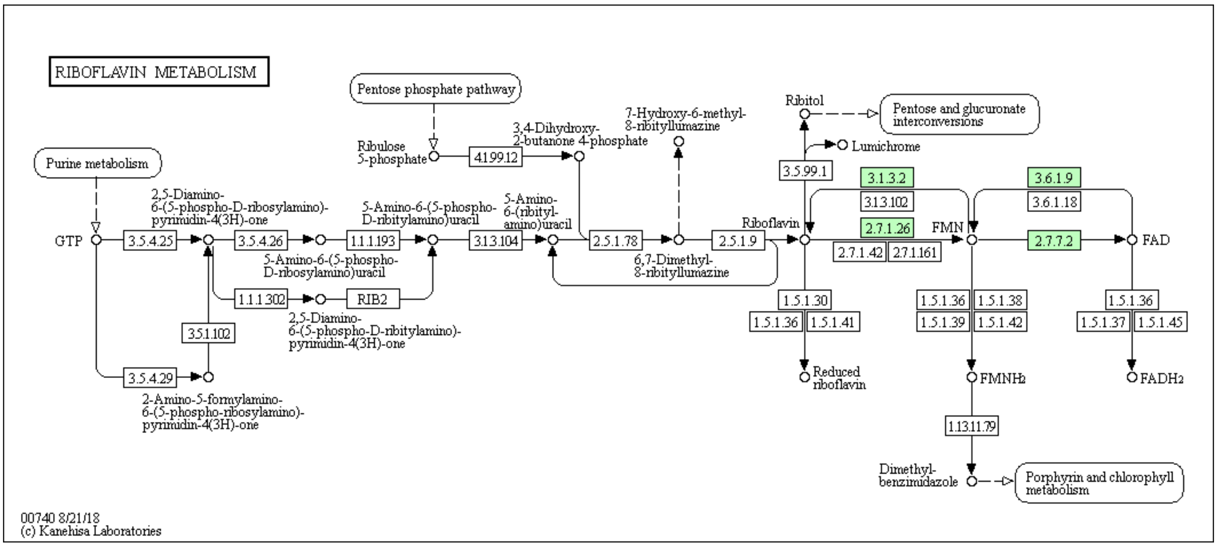


#### Supplementary Figure 9 - Riboflavin KEGG map of *Trichoplax* H1 genome

Genes present in the genome of *Trichoplax* H1 are shown in green

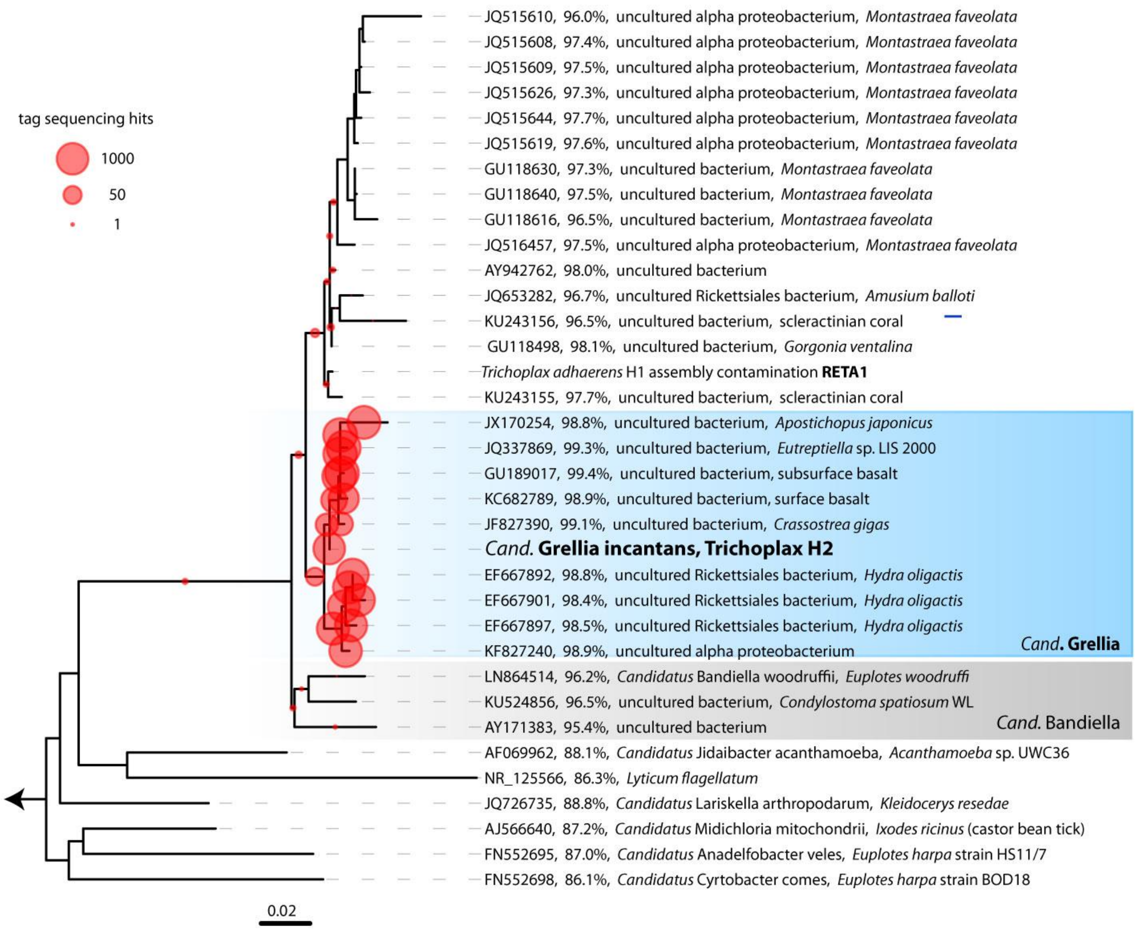


#### Supplementary Figure 10 – EPA of SRA sequences that were ≥ 99% identical to Grellia incantans in 16S rRNA gene tree of Midichloriaceae

Scale bar indicates substitutions per site. 16S rRNA tree of G. incantans and related Midichloriaceae; for each sequence, the accession number, the % identity to G. incantans, and the published taxonomic names and hosts, where available, are indicated. EPA placements of SRA sequences detected with IMNGS to be at least 99% identical to the G. incantans sequence are indicated in red circles, circle size indicates numbers of placements.

### Supplementary Tables

#### Supplementary Table 1 – The microbiome is dominated by Grellia incantans and Ruthmannia eludens.

Shown are SSU hit read counts in reads per million SSU reads that could be mapped to any of the taxa found with phyloFlash. Only taxa above 1000 SSU reads per million are shown.

| Taxon | SSUpm TR1 | SSUpm TR2 | SSUpm TR3 | SSUpm MGA | SSUpm MGB | AVERAGE |
|---|---|---|---|---|---|---|
| Trichoplax H2 | 978747 | 928343 | 941933 | 539873 | 258020 | 729383 |
| Grellia incantans | 8195 | 25212 | 19241 | 22190 | 10179 | 17003 |
| Ruthmannia eludens | 7959 | 19472 | 21559 | 5684 | 3923 | 11719 |
| Alcanivorax | 445 | 8331 | 2792 | 1347 | 19903 | 6564 |
| Marine Gammas E01-9C-26 | 0 | 8 | 3 | 10084 | 8477 | 3715 |
| unc. Alpha | 3 | 6 | 75 | 2782 | 15440 | 3661 |
| Marinobacter | 4 | 93 | 227 | 2057 | 15386 | 3553 |
| Alcanivorax | 162 | 3541 | 3040 | 0 | 5916 | 2532 |
| Erythrobacter | 10 | 52 | 67 | 7750 | 4203 | 2416 |
| Methylobacterium | 771 | 4619 | 2495 | 350 | 1652 | 1977 |
| Labrenzia | 122 | 607 | 2668 | 0 | 3578 | 1395 |

#### Supplementary Table 2 - Overview of FISH probes used in this study

| Name | Sequence (5'-->3') | Length | Formamide concentration | Fluorochrome | Reference | RDP hits - 0 mismatches | RDP hits - 1 mismatches | RDP hits - 2 mismatches |
|---|---|---|---|---|---|---|---|---|
| RUTH-1155-22 | TTTCCACAGGCAGTCTCTTGTG | 22 | 35% | Atto-647 | This study | 6 / 5x ZB3, 1x unclassified | 18 / 10x ZB3, 7x unclassified, 1x Proteobacteria | 48 / 30x ZB3, 15x unclassified, 2x Deinococcus-Thermus, 1x Proteobacteria |
| RUEL-846-22 | ATTATCTGGGGCACTGAAAGGG | 22 | 35% | Atto-594<br>4x Atto-594 | This study | none | none | 28 / 11x ZB3, 17x unclassified |
| GREL-1031-20 | CTGTGATAGTCCAGCCGAAC | 20 | 35% | Atto-647 | This study | 31/ 21 Proteobacteria, 3x Verrucomicrobia, 7x unclassified | 195 / 111x Proteobacteria, 53x Verrucomicrobia, 24x Actinobacteria, 7x unclassified | 1158 / 840x Proteobacteria, 213x Verrucomicrobia, 75x Actinobacteria, 24x unclassified, 3x Cyanobacteria, 1x Bacteroidetes, 1x Marinimicrobia, 1x Nitrospinae |
| GRIN-62-21 | GCACAAATATCGTCCGTTCGA | 21 | 35% | 4xAtto-594<br>2xAtto-647 | This study | 6 / 6x Proteobacteria | 17/ 17x Proteobacteria | 30 / 23x Proteobacteria, 6x Firmicutes, 1x Actinobacteria |
| EUB I 338 | GCTGCCTCCCGTAGGAGT | 18 | 35% | Atto-488 | [21] | - | - | - |
| EUB II 338 | GCAGCCACCCGTAGGTGT | 18 | 35% | Atto-488 | [22] | - | - | - |
| EUB III 338 | GCTGCCACCCGTAGGTGT | 18 | 35% | Atto-488 | [22] | - | - | - |
| NON 338 | ACTCCTACGGGAGGCAGC | 18 | 35% | Cy3 | [23] | - | - | - |

NOTE: Supplementary Tables 3 – 7 are available as separate supplementary files at https://doi.org/10.5281/zenodo.2583677

##### Supplementary Table 3 – Ruthmannia eludens transcriptome

##### Supplementary Table 4 – *Trichoplax* H2 transcriptome analysis

##### Supplementary Table 5 – Grellia incantans transcriptome

##### Supplementary Table 6 – Grellia incantans proteome

##### Supplementary Table 7 – Tag sequencing libraries with hits from Midichloriaceae.

#### Supplementary Videos

##### Supplementary Video 1 – Rendering of 3D reconstruction

The Supplementary video is available at https://figshare.com/s/b05fa0af88142f696476 (doi 10.6084/m9.figshare.7485683)

#### Supplementary Datasets

##### Supplementary Dataset 1 – Aligned tomography stack

The aligned tomography slices used for the reconstruction shown in Figure 4 are available at https://figshare.com/s/886b869a9ada0264ffb2 (doi 10.6084/m9.figshare.7429793).
